# Synergistic and Antagonistic Drug Interactions in the Treatment of Systemic Fungal Infections

**DOI:** 10.1101/843540

**Authors:** Morgan A. Wambaugh, Steven T. Denham, Brianna Brammer, Miekan Stonhill, Jessica C. S. Brown

## Abstract

Invasive fungal infections cause 1.6 million deaths annually, primarily in immunocompromised individuals. Mortality rates are as high as 90% due to limited number of efficacious drugs and poor drug availability. The azole class antifungal, fluconazole, is widely available and has multi-species activity but only inhibits fungal cell growth instead of killing fungal cells, necessitating long treatments. To improve fluconazole treatments, we used our novel high-throughput method, the overlap^2^ method (O2M), to identify drugs that interact with fluconazole, either increasing or decreasing efficacy. Although serendipitous identification of these interactions is rare, O2M allows us to screen molecules five times faster than testing combinations individually and greatly enriches for interactors. We identified 40 molecules that act synergistically (amplify activity) and 19 molecules that act antagonistically (decrease efficacy) when combined with fluconazole. We found that critical frontline beta-lactam antibiotics antagonize fluconazole activity. A promising fluconazole-synergizing anticholinergic drug, dicyclomine, increases fungal cell permeability and inhibits nutrient intake when combined with fluconazole. *In vivo*, this combination doubled the time-to-endpoint of mice with disseminated *Cryptococcus neoformans* infections. Thus, our ability to rapidly identify synergistic and antagonistic drug interactions can potentially alter the patient outcomes.

## Introduction

Invasive fungal infections are an increasing problem worldwide, contributing to 1.6 million deaths annually (Almeida et al., 2019; Bongomin et al., 2017; Brown et al., 2012). These problematic infections are difficult to treat for many reasons. Delayed diagnoses, the paucity and toxicity of antifungal drugs, and the already immunocompromised state of many patients result in mortality rates of up to 90% (Brown et al., 2012; Pianalto and Alspaugh, 2016; Scorzoni et al., 2017). To date, there are only four classes of antifungals, which primarily target the fungal cell envelope (cell wall and plasma membrane) (Coelho and Casadevall, 2016; Odds et al., 2003; Pianalto and Alspaugh, 2016; Scorzoni et al., 2017). The population of immunocompromised individuals is growing due to medical advancements, such as immunosuppression for transplants and chemotherapy. Emerging fungal pathogens are simultaneously increasing in both clinical burden and the number of causal species due to human activity such as agricultural drug use (Berger et al., 2017) and global warming (Almeida et al., 2019; Garcia-Solache and Casadevall, 2010). Thus, the need for more and better antifungal therapeutics is evident.

Among the most common invasive mycoses is cryptococcosis, which causes 220,000 cases and 180,000 deaths per year (Rajasingham et al., 2017). *Cryptococcus neoformans* and *Cryptococcus gattii* are the etiological agents of cryptococcosis, though nearly 95% of cases are caused by *C. neoformans* (Brown et al., 2012; Maziarz and Perfect, 2016). As *C. neoformans* is globally distributed throughout the environment, most individuals are exposed by two years of age (Goldman et al., 2001). However, systemic disease primarily occurs in the immunocompromised, particularly those with decreased T helper-1 cell reactions (Maziarz and Perfect, 2016). Accordingly, HIV/AIDS patients account for 80% of cryptococcal cases (Maziarz and Perfect, 2016; Rajasingham et al., 2017).

The primary treatment for cryptococcosis involves three different classes of antifungals. Standard care is a combination of amphotericin B (polyene class) and 5-fluorocytosine (5-FC; pyrimidine analog) for two weeks, followed by high dose azole treatment (e.g. fluconazole (FLZ)) for at least 8 weeks, and finally a low dose oral FLZ for at least 6 months (Cox and Perfect, 2018; Mourad and Perfect, 2018). Despite this, mortality rates remain as high as 80% for cryptococcal meningitis (Rajasingham et al., 2017). This is mainly due to the difficulty of obtaining ideal treatment standards. 5-FC is unavailable in 78% of countries, mostly due to licensing issues (Kneale et al., 2016; Mourad and Perfect, 2018). Without the inclusion of 5-FC in the treatment regiment, mortality increases by up to 25% (Kneale et al., 2016). Amphotericin B is administered intravenously, so treatment requires hospitalization, which is particularly challenging in areas such as sub-Saharan Africa, which has the highest burden of cryptococcal disease (Rajasingham et al., 2017). Due to these therapeutic hurdles, many patients are treated with FLZ alone, which decreases survival rates from 75% to 30% in high burden areas (Kneale et al., 2016). Additional treatment options are thus needed to prevent these unnecessary deaths.

One theoretical approach to improve treatment is synergistic combination therapy. Synergistic interactions occur when the combined effect of two drugs is greater than the sum of each drug’s individual activity (Cokol et al., 2011; Kalan and Wright, 2011). This is a powerful treatment option which has been utilized for a variety of infections (Kalan and Wright, 2011; Robbins et al., 2015; Spitzer et al., 2011; Zheng et al., 2018). Amphotericin B and 5-FC act synergistically, and mortality rates increase dramatically when one is unavailable (Beggs, 1986; Kneale et al., 2016; Schwarz et al., 2006). Synergistic interactions can also cause fungistatic drugs to switch to fungicidal, providing a more effective treatment option (Cowen et al., 2009).

Additionally, molecules can interact antagonistically to decrease therapeutic efficacy (Caesar and Cech, 2019; Roberts and Gibbs, 2018). Antagonistic interactions further complicate this already challenging infection (Khandeparkar and Rataboli, 2017; Vadlapatla et al., 2014), since immunocompromised patients are frequently treated with multiple drugs. 56% of AIDS patients experience polypharmacy, or greater than five medications (Siefried et al., 2018). Polypharmacy doubles the risk of antiviral therapy nonadherence to 49% of HIV^+^ patients (Lohman et al., 2018) and increases mortality by 68% in HIV^+^ and 99% in HIV^-^ patients (Cantudo-Cuenca et al., 2014). Better understanding of the molecular mechanisms underlying both synergistic and antagonistic drug interactions will allow us to improve identification and selection for or against these interactions.

The overlap^2^ method (O2M) uses at least one known synergistic drug pair and a large scale chemical-genetics dataset to predict synergistic and antagonistic drug interactions rapidly and on large scales (Brown et al., 2014; Wambaugh and Brown, 2018; Wambaugh et al., 2017). Each molecule of the synergistic pair induces a growth phenotype in a precise set of mutants (enhanced or reduced growth). Since these mutants exhibit the same phenotype in the presence of both molecules in a synergistic pair, we hypothesize that any other molecule eliciting the same phenotype in those mutants will also synergize with each molecule in the original pair. This method can be used against multiple microbes and applied to any published chemical-genetics dataset (Brown et al., 2014; Wambaugh et al., 2017).

In this study, we utilized previously identified synergy prediction mutants (Brown et al., 2014) to screen a library of small molecules enriched for Federal Drug Administration (FDA-)- approved molecules. We non-discriminately identified 59 molecules that interact with FLZ, either synergistically or antagonistically. When validating these new combinations, we found that even though the analysis used a *C. neoformans* dataset (Brown et al., 2014), our synergistic and antagonistic combinations acted against pathogenic fungi from multiple phyla. These include *C. deuterogattii*, *Candida* species, and multiple clinical and environmental strains of *C. neoformans*, as well as clinical isolates of the increasingly problematic and multi-drug resistant species *Candida auris* (Chowdhary et al., 2017). Furthermore, we elucidated molecular mechanisms underlying the interaction with FLZ for a few of our most clinically relevant combinations. We also demonstrate these effects in an *in vivo* model of cryptococcosis. A particularly promising synergistic combination, dicyclomine hydrochloride and FLZ, almost doubled time-to-endpoint in a murine infection model. In sum, our high-throughput method, O2M, identifies FLZ interacting molecules with potential clinical impacts.

## Results

### Synergy prediction mutants for fluconazole allow for high-throughput screening of small molecule interactions

We previously demonstrated that O2M identifies genes whose knockout mutants, termed synergy prediction mutants, exhibit phenotypes that are indicative of synergistic interactions between small molecules (Brown et al., 2014; Wambaugh et al., 2017). O2M requires a chemical-genetics dataset, in which a library of knockout mutants is grown in the presence of >100 small molecules. We calculated quantitative growth scores (slower or faster growth) for each mutant/molecule combination. This produces the “chemical genetic signature” for each molecule in the dataset. We then used these “signatures” from a known synergistic combination to identify additional combinations, the rationale being that similarities between chemical-genetic signatures of known synergistic pairs contain information that is indicative of the interaction. When we compare the chemical-genetic signatures of a pair of small molecules already known to act synergistically, we identify a subset of mutants with similar growth scores and term them “synergy prediction mutants” (Fig. 1A). We hypothesize that any molecule eliciting the same growth phenotype in these mutants would also act synergistically with either molecule in the known synergistic pair. This was completed and tested in our previous publication (Brown et al., 2014) using FLZ and its known synergistic interacting partners fenpropimorph and sertraline (Jansen et al., 2009; Zhai et al., 2012).

**Figure 1.**
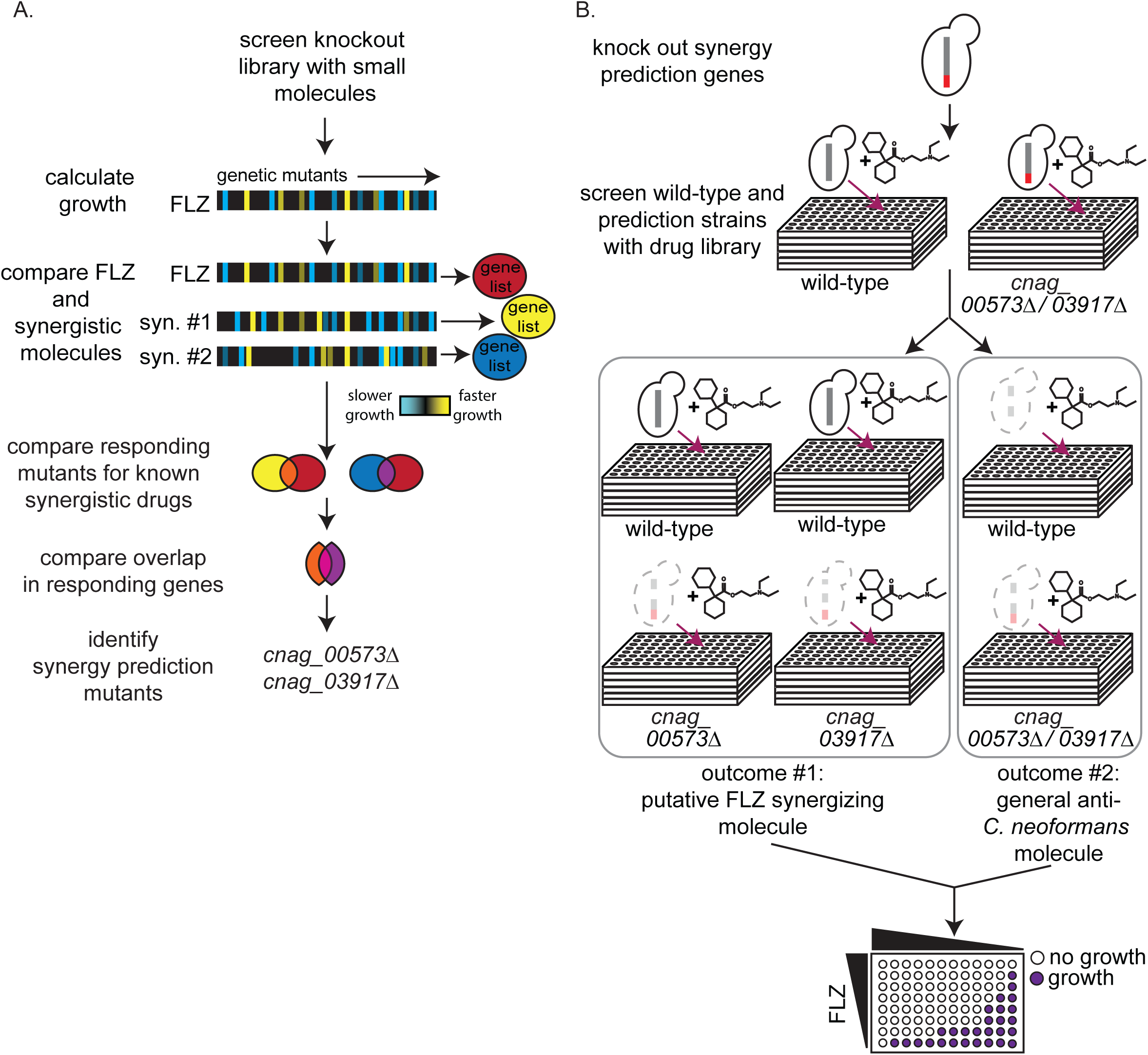
High-throughput screening for fluconazole interacting molecules using synergy prediction mutants. **A)** Outline of overlap^2^ method (O2M), which is also presented in Brown et al. and Wambaugh et al. O2M requires a chemical-genetic dataset which can be generated by growing a collection of mutants in the presence of > 100 small molecules individually. Growth scores are then calculated for each small molecule + mutant combination. In the heatmaps, the vertical line represents a different mutant. Blue represents slower growth compared to wild-type cells and yellow represents faster growth compared to wild-type cells. Comparing our starting drug (FLZ) and known synergistic molecules, we can identify genes whose knockout mutants show similar growth scores to the starting drug and all known synergistic partners. These are the synergy prediction mutants. **B)** Screening method to identify molecules that synergize with FLZ as well as anti-*C. neoformans* molecules. These molecules are then validated in a checkerboard assay.

O2M identified three gene deletion mutants (*cnag_00573*Δ, *cnag_03664*Δ, and *cnag_03917*Δ) as synergy prediction mutants for identifying interactions with FLZ (Brown et al., 2014). Using these gene mutants, we performed a high-throughput screen for synergistic interactions. Our assay is simple: differential growth between wild-type and synergy prediction mutants is indicative of a synergistic interaction with FLZ or any other starting drug. It does not require multi-drug assays, as the “synergy prediction mutant” substitutes for one of the small molecules in the interaction, phenocopying the FLZ-small molecule interaction to produce synthetic lethality. We screened the Microsource Spectrum Collection, a small-molecule library of 2,000 compounds enriched for FDA-approved molecules. We grew *C. neoformans* wild-type and synergy prediction mutants (*cnag_00573*Δ and *cnag_03917*Δ) in the presence of each small molecule (1 µM), identifying those that caused a significant difference in growth between the wild-type and both synergy prediction mutants after 48 hours of growth (Fig. 1B). The mutant *cnag_03664*Δ was not used due to its inherent slow growth. Using these synergy prediction mutants, we identified 313 putative FLZ synergistic molecules (Table S1).

We validated potential synergistic interactions in checkerboard assays, for which serial dilutions of each drug are crossed in a 96-well plate (Fig. 1B). Synergistic interactions are defined as a ≥ 4-fold decrease in the minimum inhibitory concentration (MIC) of each small molecule in the pair, resulting in a fractional inhibitory concentration index (FICI) of ≤ 0.5 (Johnson et al., 2004; Odds, 2003). We tested the 129 molecules with single agent efficacy against *C. neoformans* growth in the preferred checkerboard assay. We found that 40 molecules were synergistic with FLZ, meaning 31% of these molecules were correctly predicted by O2M (Fig. 2A). However, checkerboard assays require that both small molecules in the pair are able to inhibit growth of *C. neoformans* individually, which was not the case with all our putative synergistic molecules. In those cases, we performed Bliss Independence, which identifies whether molecules enhance the action of FLZ (Tang et al., 2015). In a 96-well plate, we created a gradient of FLZ combined with 10 µM and 100 nM concentrations of the 55 small molecules that could not inhibit *C. neoformans* alone. We found 6 of these molecules enhanced the action of FLZ at both 10 µM and 100 nM concentrations and were deemed synergistic (Fig. S1). The FLZ-synergistic molecules belonged to a wide range of bioactive categories including antidepressants, adrenergic agonists, as well as antiinfectives (Fig. 2B and Table 1).

**Figure 2.**
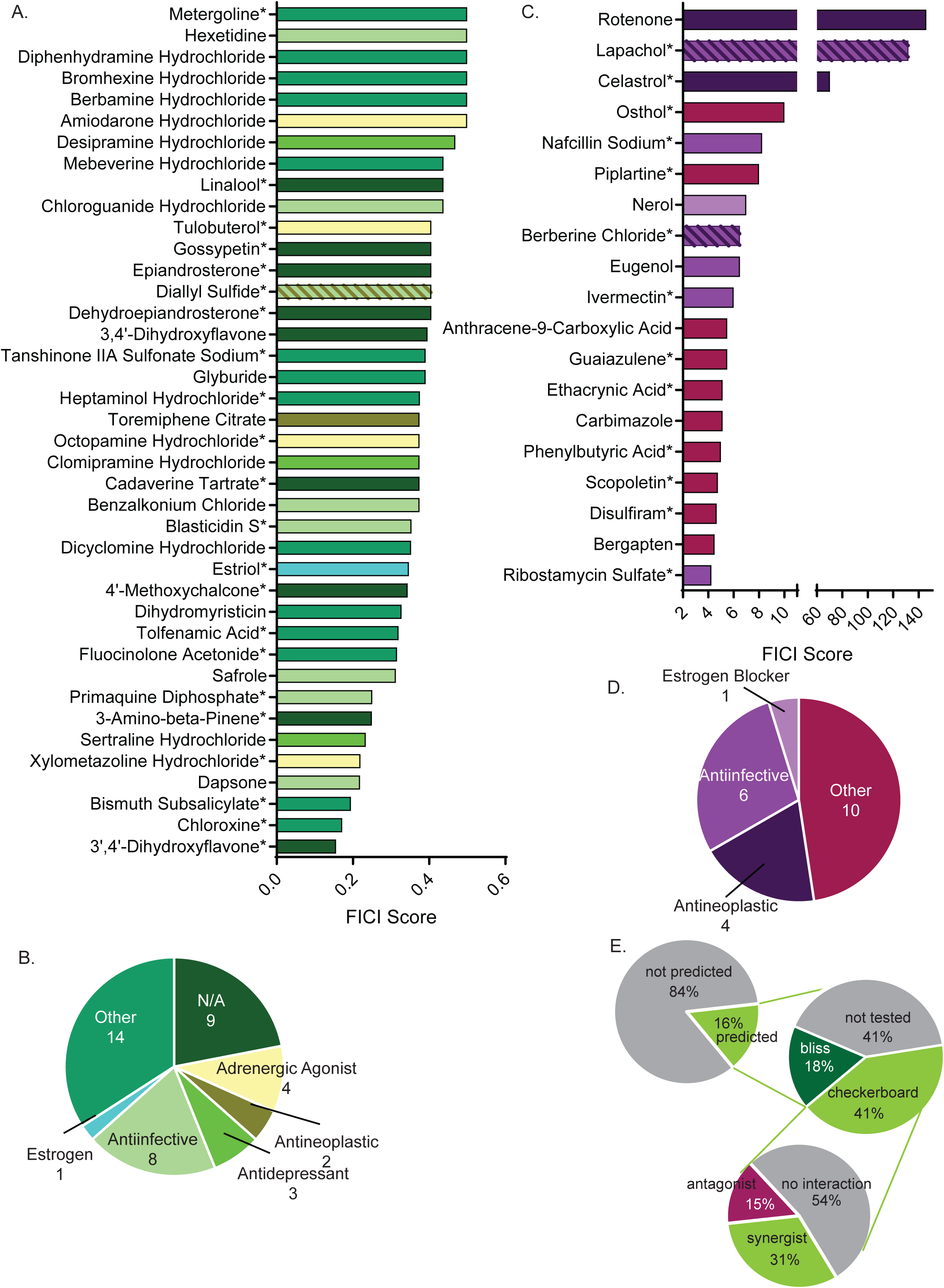
Synergistic and antagonistic molecules identified from high-throughput screen. **A)** Fractional inhibitory concentration index (FICI) score of synergistic molecules identified from our high-throughput screen. Color of bar corresponds with bioactivities listed in C. **B)** Categories of bioactivities of synergistic molecules with the corresponding number of molecules in each category. **C)** FICI scores of antagonistic molecules from screen. Colors correspond with bioactivities listed in D. **D)** Categories of bioactivities of antagonistic molecules with corresponding number of molecules. All bioactivities came from Microsource Spectrum molecule list which is also seen in Table 1. **E)** Representation of percentage of molecules predicted from the entire library (top), molecules tested in various assays (middle), and molecules yielding an interaction from checkerboards (bottom). * represents FICI for 50%inhibition of *C. neoformans* all other scores listed are the FICI for 90% inhibition (FICI90).

**Table 1.** Minimum inhibitory concentrations for fluconazole interacting molecules. All values are against *C. neoformans* strain CM18. MIC for 90% inhibition (MIC90) listed when possible. MIC50 = MIC for 50% inhibition.

| Small Molecule | Bioactivity | MIC 50 | MIC 90 | Result |
| --- | --- | --- | --- | --- |
| 3,4'-Dihydroxyflavone | N/A | 1.40 mM | 8.33 mM | synergistic |
| 3',4'-Dihydroxyflavone | N/A | 8.93 mM | N/A | synergistic |
| 3-Amino-beta-Pinene | N/A | 11.7 mM | 50.1 mM | synergistic |
| 4'-Methoxychalcone | N/A | 94.2 μM | N/A | synergistic |
| Amiodarone Hydrochloride | adrenergic agonist, coronary vasodilator, Ca channel blocker | 2.61 μM | 6.98 μM | synergistic |
| Anthracene-9-Carboxylic Acid | Cl transport inhibitor | 0.182 mM | 1.67 mM | antagonistic |
| Benzalkonium Chloride | antiinfective (topical) | N/A | 14.5 μM | synergistic |
| Berbamine Hydrochloride | antihypertensive, skeletal muscle relaxant | N/A | 63.5 μM | synergistic |
| Berberine Chloride | antiarrhythmic, α2 agonist, cholinesterase, anticonvulsant, antiinflammatory, antibacterial, antifungal, antitrypanosomal, antineoplastic, immunostimulant | 1.43 mM | 2.61 mM | antagonistic |
| Bergapten | antipsoriatic, antiinflammatory | 1.18 mM | 2.36 mM | antagonistic |
| Bismuth Subsalicylate | antidiarrheal, antacid, antiulcer | 1.49 mM | N/A | synergistic |
| Blasticidin S | antibiotic, antifungal; LD50 (rat po) 16mg/kg | 0.230 mM | 0.717 mM | synergistic |
| Bromhexine Hydrochloride | expectorant | N/A | 15.1 mM | synergistic |
| Cadaverine Tartrate | N/A | 2.27 mM | 5.82 mM | synergistic |
| Carbimazole | antithyroid | 0.215 mM | 0.859 mM | antagonistic |
| Celastrol | antineoplastic, antiinflammatory, NO synthesis inhibitor, chaperone stimulant | 68.6 μM | 0.229 mM | antagonistic |
| Chloroguanide Hydrochloride | antimalarial | N/A | 1.03 mM | synergistic |
| Chloroxine | chelating agent | 2.02 μM | 6.91 μM | synergistic |
| Clomipramine Hydrochloride | antidepressant | 0.650 mM | 0.780 mM | synergistic |
| Dapsone | antibacterial, leprostatic, dermatitis herpetiformis suppressant | 29.2 mM | N/A | synergistic |
| Dehydroepiandrosterone | N/A | 1.50 mM | 12.0 mM | synergistic |
| Desipramine Hydrochloride | antidepressant | N/A | 1.89 mM | synergistic |
| Diallyl Sulfide | antibacterial, antifungal, antineoplastic, antihypercholesterolaemic, hepatoprotectant | 7.18 mM | 32.9 mM | synergistic |
| Dicyclomine Hydrochloride | anticholinergic | N/A | 4.85 mM | synergistic |
| Dihydromyristicin | GSH transferase inducer | 0.283 M | N/A | synergistic |
| Diphenhydramine Hydrochloride | antihistaminic | 17.5 mM | N/A | synergistic |
| Disulfiram | alcohol antagonist | N/A | 7.38 μM | antagonistic |
| Epiandrosterone | N/A | 1.18 mM | 3.15 mM | synergistic |
| Estriol | estrogen | N/A | 24.8 mM | synergistic |
| Ethacrynic Acid | diuretic | 0.350 mM | 1.14 mM | antagonistic |
| Eugenol | analgesic (topical), antiseptic, antifungal | 2.40 mM | 14.3 mM | antagonistic |
| Fluocinolone Acetonide | glucocorticoid, antiinflammatory | 1.84 mM | 3.69 mM | synergistic |
| Glyburide | antihyperglycemic | 0.248 mM | 1.99 mM | synergistic |
| Gossypetin | N/A | 1.62 mM | N/A | synergistic |
| Guaiazulene | antioxidant, inhibits lipid peroxidation inhibitor, antiinflammatory, hepatoprotectant; LD50(rat) 1550 mg/kg po | 3.47 mM | 10.8 mM | antagonistic |
| Heptaminol Hydrochloride | vasodilator | 6.16 mM | N/A | synergistic |
| Hexetidine | antifungal | 50.7 μM | 0.253 mM | synergistic |
| Ivermectin | antiparasitic | N/A | 1.91 mM | antagonistic |
| Lapachol | antineoplastic, antifungal | 36.2 mM | N/A | antagonistic |
| Linalool (+) | N/A | 0.783 mM | 3.14 mM | synergistic |
| Mebeverine Hydrochloride | muscle relaxant (smooth) | 4.44 mM | 8.88 mM | synergistic |
| Metergoline | analgesic, antipyretic | N/A | 0.160 mM | synergistic |
| Nafcillin Sodium | antibacterial | 2.37 mM | N/A | antagonistic |
| Nerol | weak estrogen receptor blocker | 0.165 mM | 0.709 mM | antagonistic |
| Octopamine Hydrochloride | adrenergic agonist | 24.6 mM | N/A | synergistic |
| Osthol | N/A | 0.490 mM | 1.80 mM | antagonistic |
| Phenylbutyric Acid | antiinflammatory, antihyperammonemic (Na salt) | 0.719 mM | 2.87 mM | antagonistic |
| Pipltartine | anti-asthma, antibronchitis | 3.30 mM | 0.188 M | antagonistic |
| Primaquine Diphosphate | antimalarial | 3.73 mM | N/A | synergistic |
| Ribostamycin Sulfate | antibacterial | 9.35 mM | 18.7 mM | antagonistic |
| Rotenone | acaricide, ectoparasiticide, antineoplastic, mitochondrial poison | N/A | 1.64 μM | antagonistic |
| Safrole | anesthetic (topical) and antiseptic, pediculicide | 20.1 mM | 41.0 mM | synergistic |
| Scopoletin | NO synthesis (inducible) inhibitor, anticoagulant | 2.89 mM | 10.8 mM | antagonistic |
| Sertraline Hydrochloride | antidepressant, 5HT uptake inhibitor | 0.209 mM | 0.313 mM | synergistic |
| Tanshinone IIA Sulfonate Sodium | free radical scavenger | N/A | 1.16 mM | synergistic |
| Tolfenamic Acid | antiinflammatory, analgesia | 0.772 mM | 7.74 mM | synergistic |
| Toremiphen Citrate | antineoplastic, anti-estrogen | N/A | 27.6 mM | synergistic |
| Tulobuterol | bronchodilator, beta adrenergic agonist | 23.2 mM | N/A | synergistic |
| Xylomethazoline Hydrochloride | adrenergic agonist, nasal decongestant | 2.44 mM | N/A | synergistic |

Additionally, our screen identified antagonistic interactions. These interactions are defined by a minimum 4-fold increase in MICs causing increased fungal growth (Cetin et al., 2013). We identified 19 antagonistic interactions with FLZ (Fig. 2C). Of note, many antagonists were documented antiinfectives, including some antifungals (Fig. 2D and Table 1). The remaining 70 molecules did not interact with FLZ and represent false positives of our screen (Fig. S2A).

Overall, the O2M screen predicted that 16% of the library’s small molecules would interact with FLZ. Of the predicted interactions, 46% were validated by checkerboard assay to truly interact with FLZ (synergistic or antagonistic) (Fig. 2E). All small molecules that interact with FLZ and inhibit fungal cell growth (i.e. were tested in checkerboard assays) are listed with their MICs in Table 1. FLZ non-interacting molecules are listed in Table S2. The remaining 129 molecules predicted to interact were not tested due to unavailability, or known toxicities that would have made them impractical treatments.

### Identification of general anti-cryptococcal molecules by O2M

During the screening process, wild-type *C. neoformans* is grown in each of the small molecules alone, allowing us to identify general anti-*C. neoformans* molecules (Fig. 1B). These molecules had MICs ranging from 16 nM to 760 µM, and were mostly listed as antifungals (Table S3). However, the phenotype of a general anti-*C. neoformans* molecule can overshadow any synthetic-lethal phenotypes in the synergy prediction mutant. Therefore, we also tested these molecules for synergistic interactions in the standard checkerboard assay. Two of the general anti-*C. neoformans* molecules, sulconazole nitrate and tacrolimus, were synergistic with FLZ (Fig. S2B).

### Fluconazole-synergizing and -antagonizing responses are conserved across fungal species

We next focused on several promising synergistic molecules based on either low MICs or interesting bioactivity, with the intent of testing them against additional *C. neoformans* strains and other medically important fungi. We also selected potentially important antagonistic interactions based on bioactivities relevant to cryptococcosis patient populations. We tested these interactions against the *C. neoformans* lab strain KN99 and 10 additional environmental or clinical *C. neoformans* isolates (Chen et al., 2015). We also tested our combinations against *Cryptococcus deuterogattii*, *Candida albicans*, *Candida glabrata*, and two strains of *Candida auris*. Our interacting small molecules displayed similar MICs across these different *C. neoformans* strains and fungal species (Table S4). Of the FLZ-interacting pairs, benzalkonium chloride, berbamine hydrochloride (HCl), clomipramine HCl, dicyclomine HCl, and sertraline HCl synergistically inhibited the growth of most of the strains/species (Fig. 3B, C, E, G, J).

**Figure 3.**
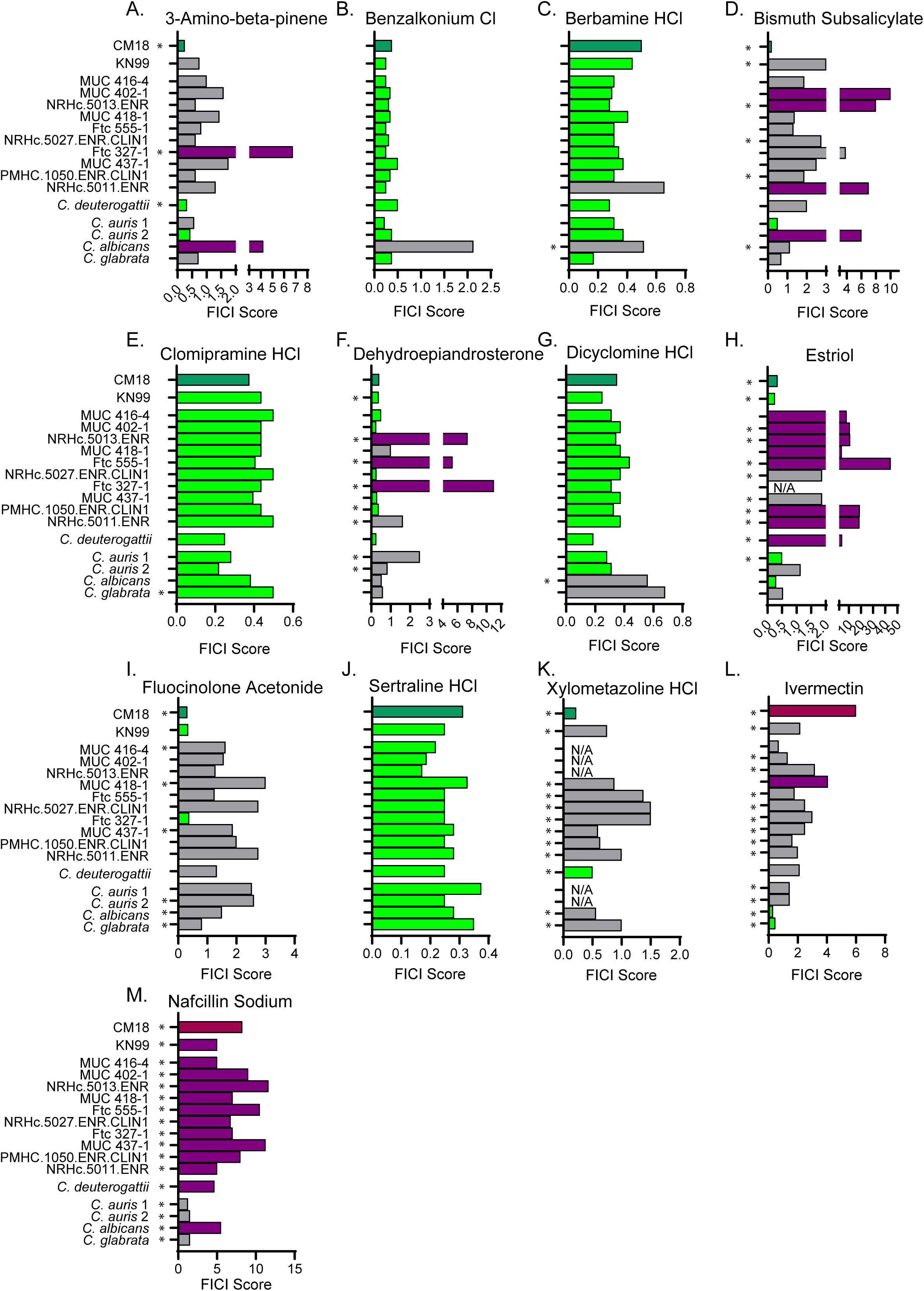
Synergistic and antagonistic combinations affect other fungal strains and species. Fractional inhibitory concentration index (FICI) scores of synergistic and antagonistic combinations with FLZ in other fungal strains/species for **A)** 3-Amino-beta-pinene **B)** Benzalkonium Cl **C)** Berbamine HCl **D)** Bismuth Subsalicylate **E)** Clomipramine HCl **F)** Dehydroepiandrosterone **G)** Dicyclomine **H)** Estriol **I)** Fluocinolone Acetonide **J)** Sertraline HCl **K)** Xylometazoline HCl **L)** Ivermectin **M)** Nafcillin Sodium. * represents FICI for 50% inhibition all other scores listed are the FICI90. Strains/species listed on left. CM18 (top) represents original result. Green bars represent FICI scores ≤ 0.5 yielding a synergistic result. Violet bars represent FICI scores ≥ 4 yielding an antagonistic result. No interactions are in grey bars. HCl = hydrochloride, Cl = chloride.

Additionally, nafcillin sodium was antagonistic with FLZ in most of the strains/species (Fig. 3M). The other synergistic or antagonistic molecule pairs had more variable results among the different strains/species (Fig. 3), highlighting the importance of testing putative antimicrobials against multiple strains when considering therapeutic relevance.

### Molecular structure predicts additional synergistic interactions

Upon identifying new FLZ-based synergistic pairs, we sought to predict other FLZ synergizing molecules based on structure alone. We chose four of our newly identified synergistic molecules that contained large ring structures (a commonality amongst our synergists): dicyclomine HCl, desipramine HCl, sertraline HCl, and diphenhydramine HCl. We used both ChemSpider (Pence and Williams, 2010) and ChemMine MSC Similarity Tool to identify structurally similar molecules (Backman et al., 2011). We chose three molecules (proadifen, drofenine, and naftidrofuryl) with ≥50% structure similarity to dicyclomine HCl, and all were synergistic with FLZ in checkerboard assays (Fig. S3A-D, M). We tested three molecules (impramine, mianserine, and lofepramine) with >40% structural similarity to desipramine, two of which were synergistic with FLZ (Fig. S3E-H, M). Lofepramine, which was not synergistic, is the prodrug to desipramine. Sibutramine, shares >40% structural similarity with sertraline, and is synergistic with FLZ as well (Fig. S3I, J, M). Lastly, we tested citalopram, which shares >45% structural similarity to diphenhydramine, but was not synergistic with FLZ (Fig. S3K-M). Overall, 75% of the structurally similar molecules proved to be synergistic with FLZ, demonstrating that structure can serve as a powerful basis to predict additional synergistic combinations prior to elucidating mechanism of action.

### Exposure to beta-lactam antibiotics increases ergosterol levels and antagonizes fluconazole activity

Among the antagonists, we were particularly interested in the interaction between nafcillin sodium (nafcillin) and FLZ that emerged in multiple fungal strains and species (Fig. 3M). Nafcillin is a common penicillinase-resistant penicillin antibiotic (Letourneau, 2019; Nathwani and Wood, 1993). Furthermore, the cryptococcosis patient population, which consists of mainly HIV/AIDS patients, is at high risk for multiple infections, increasing the likelihood that they could require overlapping treatments for bacterial and fungal infections (Kaplan et al., 2009). Thus, we first wanted to determine if other beta-lactam antibiotics would also antagonize fluconazole activity (Fig. 4A-M). Antagonism with FLZ was not a universal attribute among penicillin antibiotics, but was evident for oxacillin and methicillin (Fig. 4N). We also tested a first-generation cephalosporin, cefazolin, that is often prescribed in place of nafcillin (Letourneau, 2019; Miller et al., 2018) and a second-generation cephalosporin, cefonicid. We found that both of these molecules also antagonize FLZ (Fig. 4N). Of the beta-lactams tested, those that are often used to treat *Staphylococcus aureus* were antagonistic with FLZ (Fowler Jr. and Holland, 2018; Letourneau, 2019). Due to this, we decided to also test alternative treatment molecules for *S. aureus* infections, particularly vancomycin and linezolid (Fowler Jr. and Holland, 2018; Lowy, 2019), both of which we found to have no interaction with FLZ (Fig. 4N). To investigate the mechanism of antagonism, we looked to other known drug interactions involving nafcillin. In particular, nafcillin antagonizes warfarin and other drugs *in vivo* by inducing cytochrome P450 enzymes through an unknown mechanism, which increases warfarin metabolism (King et al., 2018; Wungwattana and Savic, 2017). FLZ inhibits a fungal cytochrome P450 enzyme, 14α-demethylase, which halts ergosterol biosynthesis and fungal growth (Fig. 4O) (Odds et al., 2003). We hypothesized that nafcillin may also induce cytochrome P450 enzymes, such as 14α-demethylase, in *C. neoformans*, counteracting FLZ’s mechanism of action. To test this hypothesis, we examined whether nafcillin affects ergosterol biosynthesis. We extracted sterols from *C. neoformans* cells grown in the presence of nafcillin, FLZ, nafcillin + FLZ, or vehicle. We found a concentration-dependent increase in ergosterol with nafcillin treatment (Fig. 4P). Ergosterol levels in nafcillin + FLZ are not statistically different from the control treatment (Fig. 4P). Finally, we tested whether nafcillin is synergistic with the antifungal amphotericin B. Since amphotericin B kills target cells by binding and extracting ergosterol from the plasma membrane (Anderson et al., 2014), we hypothesized that if nafcillin increases ergosterol levels in *C. neoformans*, nafcillin would act synergistically with amphotericin B by increasing amphotericin B binding sites (i.e. ergosterol). Indeed, amphotericin B was synergistic with nafcillin in checkerboard assays (Fig. 4Q), further suggesting that nafcillin increases ergosterol levels in *C. neoformans*.

**Figure 4.**
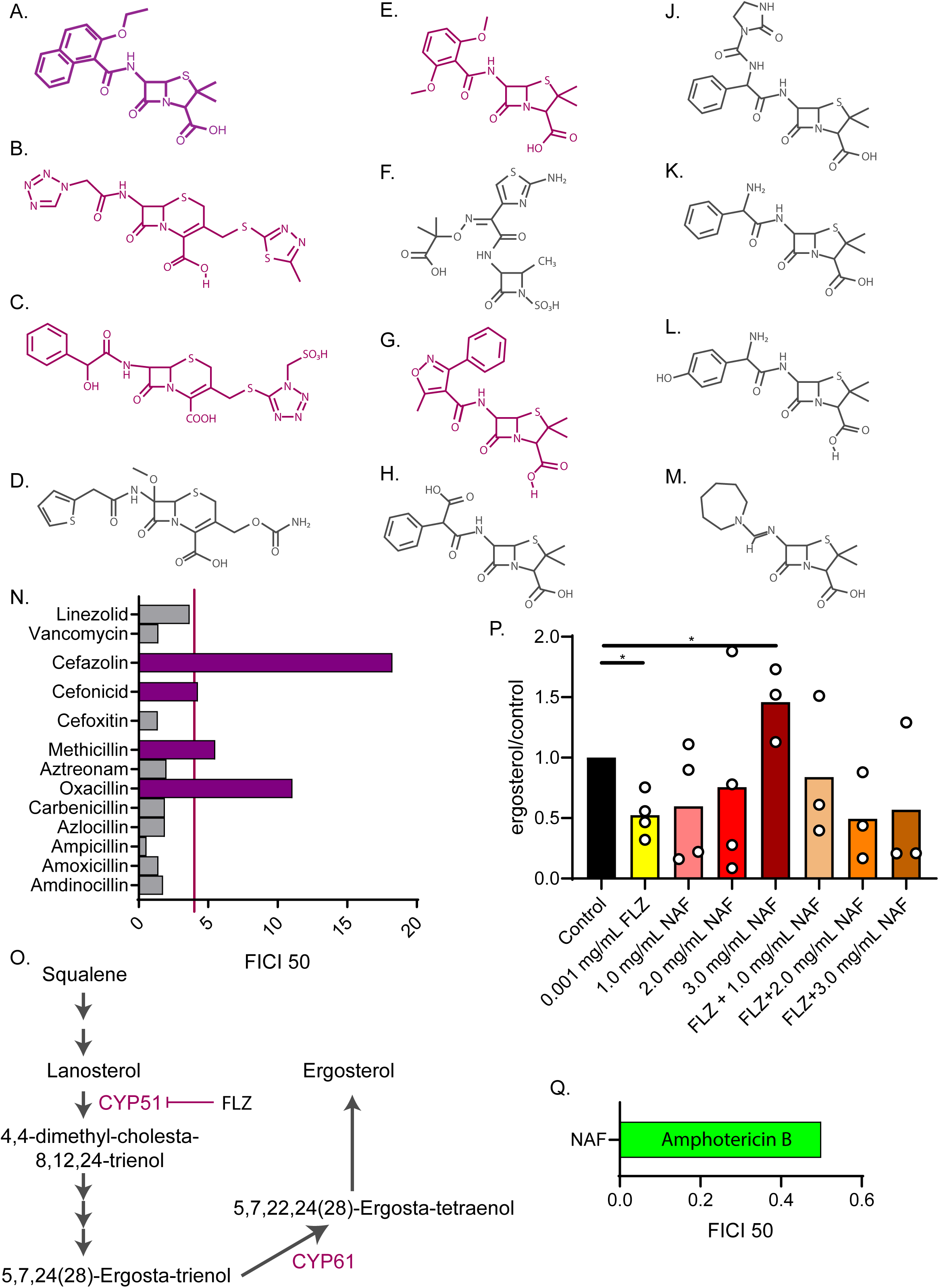
Nafcillin Sodium affects ergosterol levels. Molecular structures of beta-lactam antibiotics shown for **A)** Nafcillin Sodium **B)** Cefazolin **C)** Cefonicid **D)** Cefoxitin **E)** Methicillin **F)** Aztreonam **G)** Oxacillin **H)** Carbenicillin **J)** Azlocillin **K)** Ampicillin **L)** Amoxicillin **M)** Amdinocillin. **N)** FICI scores for 50% inhibition of *C. neoformans* of various antibiotics related to nafcillin sodium tested with fluconazole. Violet bars over the red line illustrate a FICI score of ≥ 4 indicating antagonism. **O)** Ergosterol biosynthesis pathway illustrating cytochrome P450 enzymes. **P)** Ergosterol quantification from cell treated with Nafcillin (NAF), FLZ, or NAF+FLZ. Data normalized to control treated. **Q)** FICI for 50% inhibition of *C. neoformans* treated with nafcillin sodium in combination with amphotericin B. * = p value is 0.0268 (Mann-Whitney test).

### The synergistic interaction between dicyclomine HCl and FLZ affects cell permeability and nutrient uptake

Finally, we investigated the drug dicyclomine HCl (dicyclomine), an anticholinergic agent (Table 1). This FDA-approved drug, also known as Bentyl, is used to treat urinary incontinence (Malone and Okano, 1999; Page and Dirnberger, 1981). In humans it targets a G-protein coupled receptor (GPCR) encoded by the *CHRM1* gene (Consortium, 2018; Kilbinger and Stein, 1988). *C. neoformans* does not have a *CHRM1* ortholog, but there are a large number GPCR in fungi that could be potential targets (Xue et al., 2008). When we screened a deletion library of *C. neoformans* for mutants resistant to dicyclomine, we found that 44% of the annotated dicyclomine-resistant mutants were involved in transport and trafficking, suggesting that those processes may be related to dicyclomine’s mechanism (Fig. 6A and Table S5). Thus, we hypothesized that dicyclomine alters Golgi transport. In *Saccharomyces cerevisiae*, simultaneously inhibiting Golgi trafficking and blocking ergosterol synthesis leads to mislocalization of essential plasma membrane transporters (Estrada et al., 2015). We hypothesized that the combination of dicyclomine and FLZ will phenocopy this effect in *C. neoformans* (Fig. 6B). Using propidium iodide internalization as a measure of cell permeability, we observed that dicyclomine, similar to FLZ, permeabilizes fungal cells at high doses (Fig. 6C, D, F, G). Furthermore, we saw a greater than additive increase in permeability when fungal cells were treated with low concentrations of dicyclomine and FLZ in combination (Fig. 6E and H). dicyclomine-induced permeability appeared to be independent of significant changes to cell wall chitin (Fig. S4A-F).

We next tested if dicyclomine + FLZ disrupted nutrient transporter function by measuring uptake of amino acids. If amino acid permeases are not localized to the plasma membrane, fungal cells are resistant to toxic amino acid analogs (Roberg et al., 1997). Using the same low doses of FLZ and DIC that alone do not permeabilize cells, *C. neoformans* is susceptible to the effect of either 5-fluoroanthranilic acid (5-FAA) or 5-methyl-tryptophan (5-MT). When the dose is combined, cells now show resistance (Fig. 6I-L and Fig. S4G-J). This demonstrates that certain amino acid transporters’ function is decreased and they thus may be mislocalized, conferring resistance to toxic forms of tryptophan.

### Dicyclomine + FLZ act synergistically *in vivo* and enhances survival of mice with cryptococcosis

We intranasally inoculated CD-1 outbred mice (Charles River) with *C. neoformans* and allowed the infection to progress 8 days. Colony forming unit data indicated that at this point 100% of the mice exhibited fungal dissemination to the liver, and 40% exhibited dissemination to the brain (Fig. S5). A disseminated infection is consistent with human patients at treatment onset, as patients often don’t seek treatment until *C. neoformans* has disseminated to the brain (Zhu et al., 2010). From 8 days post-inoculation (d.p.i) until 40 d.p.i., we administered dicyclomine, FLZ, dicyclomine + FLZ or PBS (vehicle) intraperitoneally. We used doses of both FLZ and dicyclomine which were within the range of doses given to humans (Lexicomp, 1978-2019a, b). We sacrificed mice when they reached 80% of their initial mass (survival endpoint). Dicyclomine at either a low or high dose did not affect mouse survival compared to PBS-treatment. However, dicyclomine in combination with FLZ significantly improved survival over FLZ alone in a dose-dependent manner (Fig. 6M), indicating that dicyclomine may not be effective at treating cryptococcosis on its own, but could well be of therapeutic benefit when combined with FLZ.

## Discussion

Synergistic combination therapies are increasingly important clinical options, especially for drug resistant microbes (Cowen and Lindquist, 2005; Kalan and Wright, 2011; Uppuluri et al., 2008; Zheng et al., 2018). Traditionally, synergistic drug pairs were discovered serendipitously, but new methods are improving our ability to uncover important interactions (Brown et al., 2014; Cokol et al., 2011; Cokol et al., 2018; Jansen et al., 2009; Robbins et al., 2015; Spitzer et al., 2011; Wambaugh et al., 2017; Wildenhain et al., 2015). In this study, we identified a wide variety of molecules that interact synergistically with the antifungal FLZ to inhibit fungal growth. We do so without the use of noisy multi-drug assays, allowing for rapid and scalable screening. We also identify and investigate antagonistic interactions, which are clinically important (Khandeparkar and Rataboli, 2017; Vadlapatla et al., 2014) but have not been investigated in a systemic manner.

Of the 59 FLZ interacting molecules we identified, 10 have been previously described in various fungi, though 3 of those were reported as synergists while our testing revealed them to be antagonists (Ahmad et al., 2010; Butts et al., 2014; Cardoso et al., 2016; Eldesouky et al., 2018; Kang et al., 2010; Li et al., 2018; Marchetti et al., 2000; Quan et al., 2006; Robbins et al., 2015; Spitzer et al., 2011; Zhai et al., 2012). Additionally, we discovered synergistic molecules with a wide variety of bioactivities, suggesting that O2M identifies synergistic molecules with molecular mechanisms that can differ from that of the starting synergistic pair. O2M analysis for FLZ synergizers used fenpropimorph and sertraline. Fenpropimorph inhibits ergosterol biosynthesis (Marcireau et al., 1990) and sertraline inhibits protein translation (Zhai et al., 2012).

Furthermore, we identified broad spectrum interactions. All the combinations tested showed efficacy against multiple clinical and environmental isolates of *C. neoformans*, as well as *C. deuterogattii*, a related species which can cause disease in apparently immunocompetent individuals (Applen Clancey et al., 2019). We also tested our combinations against common *Candida* species that often develop multi-drug resistance (Colombo et al., 2017), including the emerging MDR pathogen *Candida auris* (Fig. 3). Our data demonstrate that FLZ synergizers and antagonizers exhibit broad activities against multiple species and isolates.

Another key component of drug discovery is the ability to improve upon the efficacy of known drugs through synthetic modification and/or identification of structurally related molecules. We found that structural similarity predicts synergistic interactions (Fig. S3), just as drugs of similar structure have similar function. These data will open up our ability to rapidly identify many more synergizers and antagonizers from a single example.

We investigated the antagonistic interaction between FLZ and nafcillin, as nafcillin is a commonly used beta-lactam antibiotic for *Staphylococcus aureus* and other difficult to treat bacterial infections (Letourneau, 2019). Patients with these infections include some of the same patients at risk for cryptococcosis (HIV and cancer patients) (Kaplan et al., 2009; Utay et al., 2016) and other fungal infections, including *C. auris* (Rudramurthy et al., 2017). When we examined nafcillin-related molecules, we found that methicillin and oxacillin also antagonize FLZ. Furthermore, two cephalosporins often used in place of nafcillin, cefonicid and cefazolin, also unexpectedly antagonistize fluconazole activity. Nafcillin has previously been shown to adversely affect patients on the drug warfarin due to nafcillin’s induction of cytochrome P450 (King et al., 2018), which decreases warfarin concentration. This has been seen in a similar combination of FLZ and the antibiotic rifampicin (Panomvana Na Ayudhya et al., 2004).

Rifampicin is a potent inducer of drug metabolism due to elevation of hepatic cytochrome P450 through increased gene expression (Bolt, 2004). This combination did indeed lower the serum levels of FLZ (Panomvana Na Ayudhya et al., 2004), which resulted in relapse of cryptococcal meningitis (Coker et al., 1990). We hypothesize that an analogous process occurs when nafcillin is combined with FLZ, with nafcillin increasing ergosterol biosynthesis enzymes, counteracting FLZ’s activity. In a recent autopsy study, 10 or 16 patients who died of cryptococcosis were administered either a penicillin or a cephalosporin (Hurtado et al., 2019). We recommend that these patients receive linezolid or vancomycin instead, since these drugs are used for similar bacterial targets but do not antagonize fluconazole activity (Fig. 4N).

Our data demonstrate that O2M identifies promising new antifungal treatments that can rapidly move into the clinic. Our new combination of dicyclomine + FLZ almost doubled the median time-to-endpoint of mice treated with human dosages of dicyclomine (Fig. 5M), which is lower than dicyclomine’s fungal MIC. Dicyclomine is orally bioavailable and able to cross the blood brain barrier (Das et al., 2013; Koerselman et al., 1999), which makes it particularly promising for fungal meningitis treatment. Since dicyclomine, like many of our FLZ synergizers, is approved by the FDA for other indications, it could rapidly move into the clinic.

**Figure 5.**
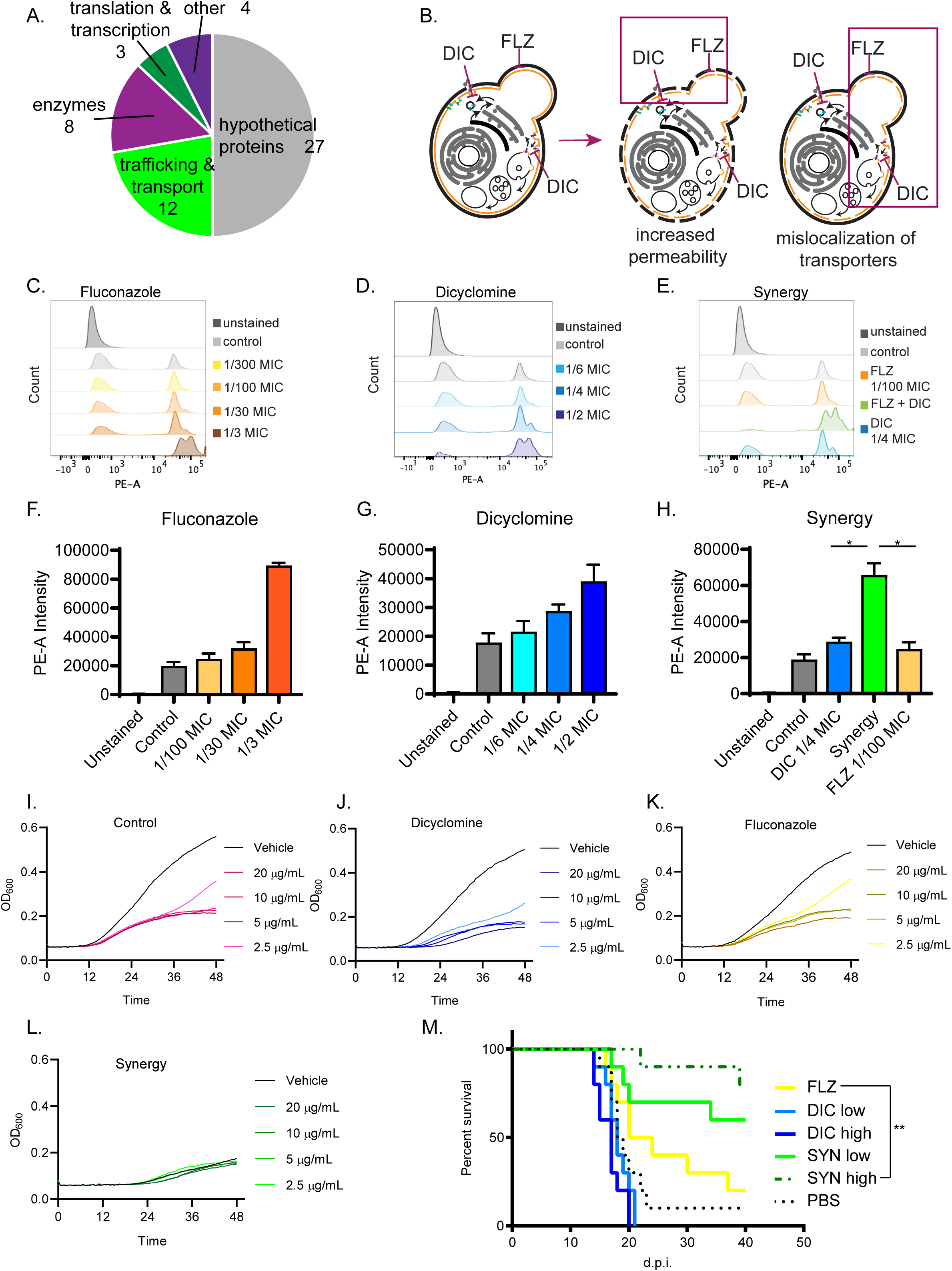
Dicyclomine affects permeability and nutrient transporters. **A)** Pie chart with processes of deletion mutants that were resistant to dicyclomine. Numbers represent number of mutants. **B)** Prediction for dicyclomine (DIC) + FLZ synergy mechanism. **C-E**) Representative flow plots of propidium iodide staining. **F-H**) Quantification of propidium iodide staining. **I-L**) Growth curves of *C. neoformans* with and without various concentrations of 5-FAA in addition to control, dicyclomine, fluconazole, or synergy treatment. **M)** Survival of CD-1 outbred mice given FLZ (8 mg/kg), DIC low (1.15 mg/kg), DIC high (2.30 mg/kg), Synergy (SYN) low (FLZ + 1.15 mg/kg DIC), SYN high (FLZ + 2.30 mg/kg) or PBS treatments. N=10. * = p value is 0.0286 (Mann-Whitney test); ** = p value is 0.0036 (Mantel-Cox test).

In sum, O2M considerably streamlined the identification of important drug interactions affecting *C. neoformans* growth. These interactions are both synergistic and antagonistic among multiple fungal species capable of causing disease in humans. We focused on FDA-approved molecules to bypass the time and considerable expense it takes to develop a new drug (Pushpakom et al., 2018). However, our method would work equally well on any library of small molecules or biologic drugs to discover new antifungals. We showed that identifying these drug interactions can quickly lead to additional interacting pairs by examining structure (Fig. S3) or by investigating underlying mechanism (Wambaugh et al., 2017). Finally, our newly discovered interaction of dicyclomine and FLZ exhibited therapeutic potential *in vivo*, demonstrating the clinical potential of fluconazole-containing synergistic pairs in the clinic.

## Acknowledgements

We thank the University of Utah Metabolomics Core for sterol quantification, Jerry Kaplan, Ph.D. for advice on sterol extraction, and members of the Brown and Mulvey labs for helpful discussion and feedback. This work was supported by NIH grant R01AI137331 and funds from the Pathology Department at the University of Utah to JCSB.

## Author Contributions

MAW and JCSB designed experiments. MAW, STD, BB, MS conducted experiments. MAW wrote the paper with input from JCSB. MAW, STD, and JCSB edited the paper.

## Methods

### Fungal strains

Screening, validation, and structurally similar assays were performed with CM18 lab strain of *C. neoformans*. Screening with synergy prediction mutants (*CNAG_00573Δ* and *CNAG_03917Δ*) was in the CM18 background. Mechanistic studies were tested using the KN99 lab strain of *C. neoformans*. Clinical and environmental isolates of *C. neoformans* tested were a gift from Dr. John R. Perfect. *C. deuterogattii* strain R265 was purchased from ATCC. *Candida albicans* reference strain SC5314 and *Candida glabrata* reference strain CBS138 were used. *Candida auris* strains AR0383 and AR0384 were from the CDC.

### Microsource Spectrum library screen

We inoculated either CM18 wild-type or *CNAG_00573Δ* or *CNAG_03917Δ* cells at 1000 cells per well of YNB + 2% glucose, then added small molecule to a final concentration of 1µM. Plates were incubated for 48 hours at 30 °C. OD_600_ was measured on the BioTek plate reader model Synergy H1. Small molecules that altered growth by absolute value 0.22 in both synergy prediction mutants was considered significant.

### *C. neoformans* growth and small-molecule assays

All assays were performed in 1x YNB + 2% glucose. To determine MICs, an overnight culture was grown at 30 °C with rotation, diluted to OD_600_ = .02925 and 1000 cells were added to each well (2 µL of culture into 100 µL of media per well). Plates were incubated at 30 °C unless otherwise stated. Small molecule gradients were diluted in 2-fold dilution series. MIC values were calculated after 48 hours of incubation.

### Checkerboard Assay and FICI calculations

We followed previously published methods (Hsieh et al., 1993; Orhan et al., 2005). Starting inoculation of either fungal strain was 2µL of an OD_600_ = 0.02925 (1000 cells per well of 100 µL medium). Plates were grown statically for 48 hours at 30 °C with minor shaking/resuspension of cells at 24 and 48 hours. Checkerboards were read at 0 and 48 hours on a BioTek plate reader model Synergy H1 (*Candida albicans* and *Candida glabrata* were read at 0, 24, and 48 hours). Growth inhibition was assessed and FICIs for 50% and 90% inhibition were determined. Repeated results were averaged for the average FICI.

### Bliss independence Assay

We created a gradient of fluconazole in a 96-well plate, then added small molecules at 10 µM or 100 nM final concentrations or vehicle. CM18 wild-type was added at 1000 cells per well of YNB+ 2% glucose. Percent growth was calculated for fluconazole, combinations, or small molecules alone. We then determined if growth inhibition caused by the combination was equal or greater than growth inhibition of the small molecules alone. The most repeatable result of multiple experiments was then averaged for average Bliss Independence Score.

### Sterol extraction

KN99 culture was grown overnight in YNB + 2% glucose. Cells were sub-cultured into various treatments (vehicle control was YNB + 2% glucose). 6 ODs of each culture were harvested and lyophilized overnight. Pellets were resuspended in 25% alcoholic potassium hydroxide, vortexed, and incubated at 85 °C water bath for 1 hour. Water and *n*-heptane were added to each tube, vortexed, and the *n*-heptane layer was transferred to borosilicate glass tubes.

### Metabolomics

Metabolomics analysis was performed at the Metabolomics Core Facility at the University of Utah which is supported by 1 S10 OD016232-01, 1 S10 OD021505-01, and 1 U54 DK110858-01.

### Resistance to dicyclomine screen

YNB + 2% glucose agar plates with or without 1.65 mg/mL of dicyclomine were made. Deletion mutants in KN99 strain were grown in YNB + 2% glucose then pinned to YNB plates. Plates were assessed at 1, 2, and 3 days for resistance.

### Cell permeability assay

An overnight culture of KN99 was grown in YNB + 2% glucose. This was then sub-cultured into the various treatments (vehicle control was either 0.1% DMSO or only YNB + 2% glucose). Cultures were grown at 30 °C with rotation for 24 hours. Cultures were washed twice and resuspended in PBS. 3 µL of propidium iodide (stock concentration = 1 mg/mL) was added to the FACS tube. After 1 min flow cytometry was performed. Voltage gates used were as follows: FSC: 500; SSC: 310; PE: 496.

### Resistance to toxic amino acids

An overnight culture of KN99 was grown in YNB + 2% glucose. This was sub-cultured into either dicyclomine (0.3 mg/mL), FLZ (3E-4 mg/mL), Synergy, or Vehicle (YNB + 2% glucose) and grown at 30 °C with rotation for 24 hours. Cells were then sub-cultured again into honeycomb plates with those previous treatments (dicylomine, FLZ, Synergy, or Vehicle) with either 20, 10, 5, 2.5 ug/mL of 5-FAA or 0.4 mg/mL 5-MT or vehicle (3.2% DMSO). Plates were read on a Bioscreen machine at 30 °C with OD_600_ measured every 30 minutes for 48 hours.

### Calcofluor white flow cytometry

An overnight culture of KN99 was grown in YNB + 2% glucose then sub-cultured into either dicylomine, FLZ, Synergy, or Vehicle (YNB + 2% glucose) and grown at 30 °C with rotation for 24 hours. Cells were washed with PBS and calcofluor white added to a final concentration of 50 µg/mL and stained for 5-15 minutes. Cells were washed once more with PBS and assessed by flow cytometry. Voltage gates as follows: FSC: 500; SSC: 317; BV421-A: 185. Significance determined with Mann-Whitney test.

### Murine model

∼8-week-old female CD-1® IGS outbred mice (Charles River Laboratories) were intranasally inoculated with *C. neoformans* strain KN99 and monitored for survival as follows. The inoculum was prepared by culturing *C. neoformans* overnight in liquid YNB+2% glucose medium. *C. neoformans* cells collected from the overnight culture were washed twice in 1XPBS, counted, and suspended at a concentration of 5×10^5^ cells / mL. Mice were anesthetized with ketamine/dexmedetomidine hydrochloride (Dexdomitor) administered intraperitoneally (i.p.) and suspended by their front incisors on a line of thread. The inoculum was delivered intranasally by pipetting 50 μL (2.5×10^4^ cells / mouse) dropwise into the nostrils. After 10 minutes, mice were removed from the thread and were administered atipamezole (Antisedan) i.p. as a reversal agent. Mice were massed daily and euthanized by CO_2_ asphyxiation when they reached 80% of their initial mass. Beginning 8 days post-inoculation, they received daily i.p. injections of either 8 mg/kg FLZ, 1.15 mg/kg DIC, 2.30 mg/kg DIC, 8 mg/kg FLZ + 1.15 mg/kg DIC, 8 mg/kg FLZ + 2.30 mg/kg DIC, or PBS control (vehicle). Dosages were determined from human doses (Lexicomp, 1978-2019a, b).

---

**Figure S1.**
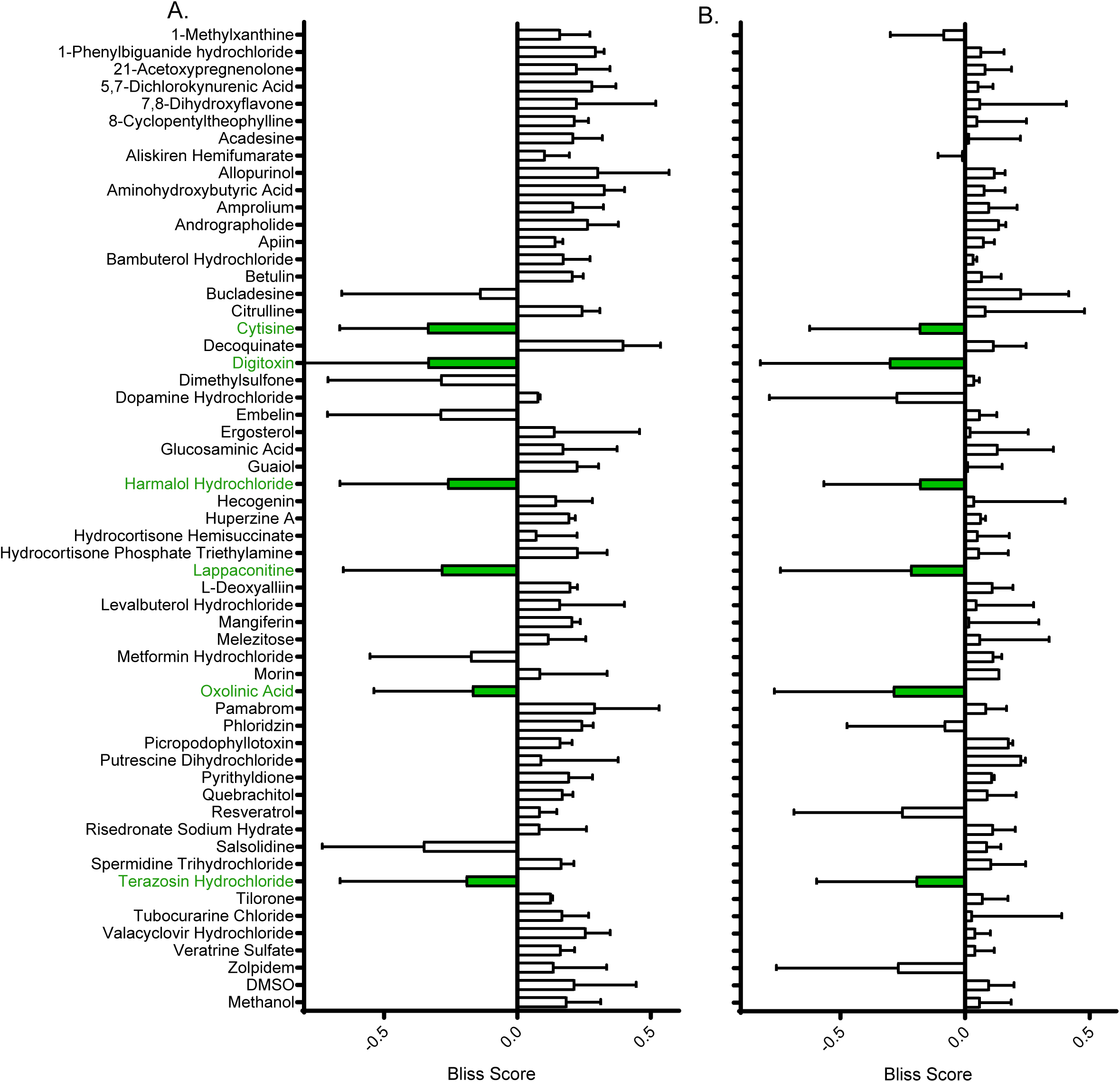
Bliss independence scores of non-single agent molecules. Average bliss independence score of molecules at 10 µM (**A**) and 100 nM (**B**). Molecules were considered synergistic if they exhibited a negative score in both concentrations the majority of times tested (green labels).

**Figure S2.**
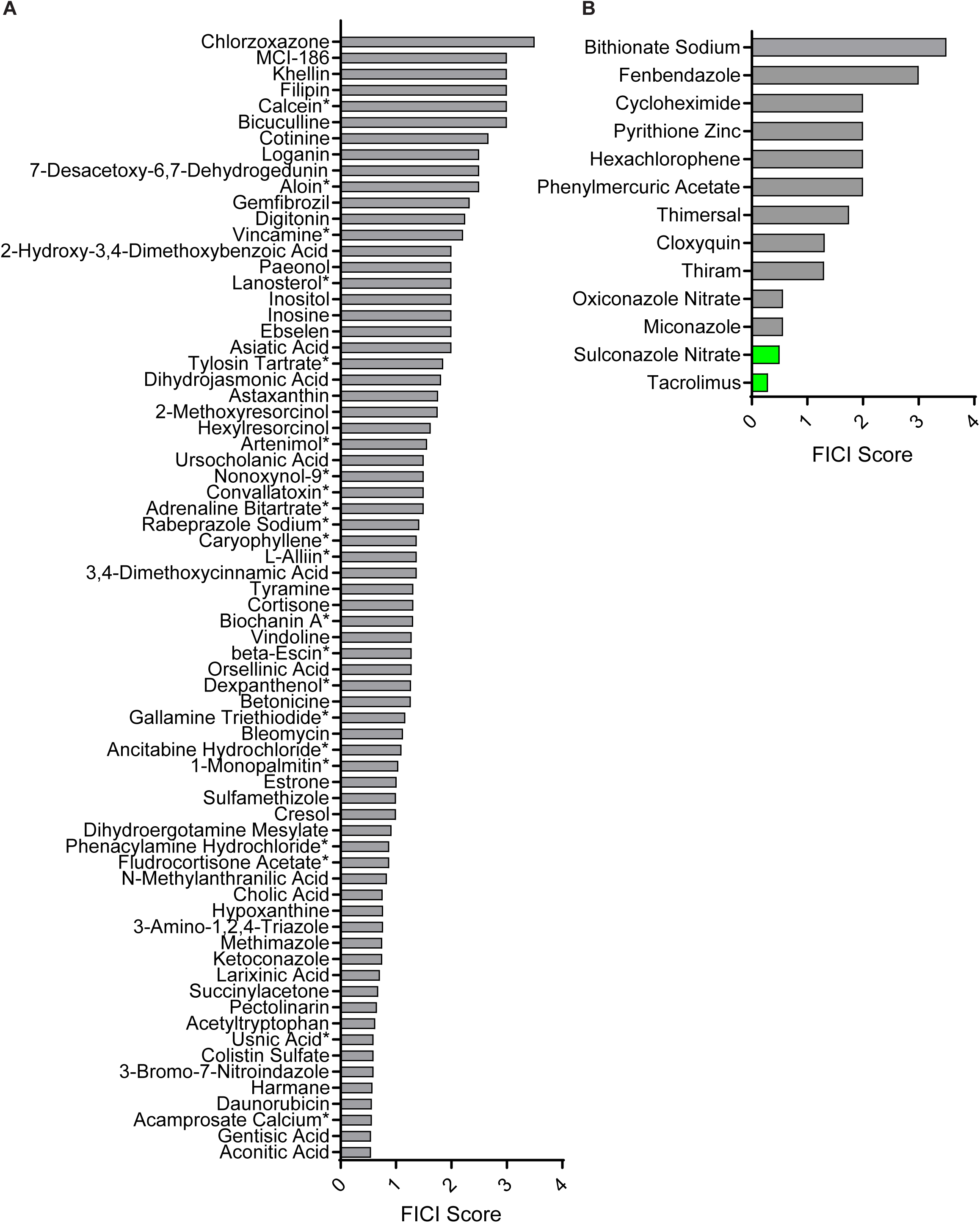
FICI scores of non-interacting molecules and general antifungals. **A**) Fractional inhibitory concentration index (FICI) of molecules identified from our high-throughput screen that did not interact with FLZ. **B**) FICI scores of molecules identified as general anti-*C. neoformans* molecules from our high-throughput screen. Synergistic interactions with FLZ labeled in green and non-interacting molecules labeled in grey. * represents FICI for 50% inhibition of *C. neoformans* all others are FICI for 90% inhibition.

**Figure S3.**
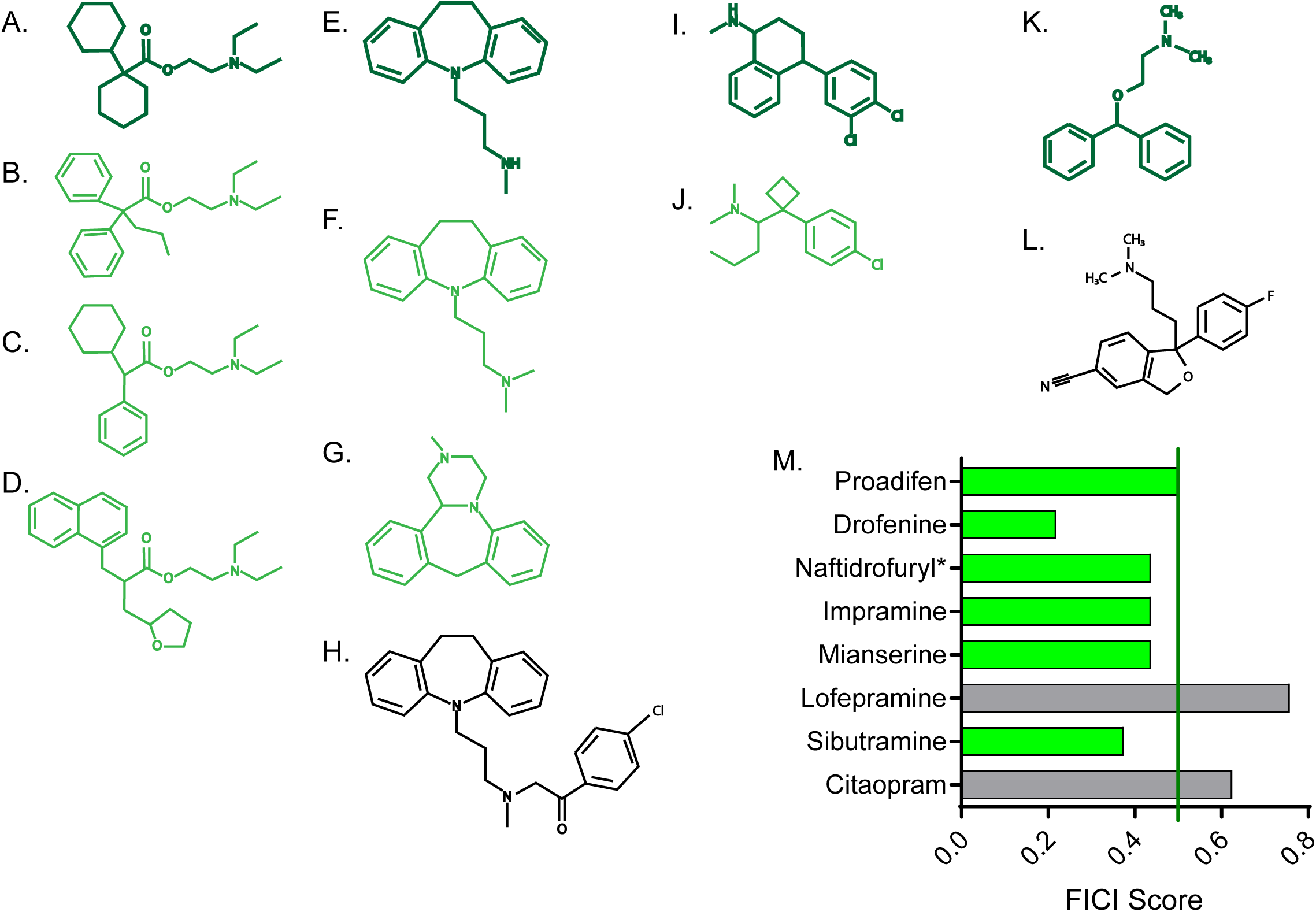
FICI scores of structurally similar molecules. Chemical structure of dicyclomine HCl (**A**) and structurally similar molecules proadifen (**B**), drofenine (**C**) and naftidrofuryl (**D**). Chemical structure of desipramine HCl (**E**) and structurally similar molecules impramine (**F**), mianserine (**G**), and lofepramine (**H**). Chemical structure of sertraline HCl (**I**) and structurally similar sibutramine (**J**). Chemical structure of diphenhydramine HCl (**K**) and structurally similar citalopram (**L**). **M**) Fractional inhibitory concentration index (FICI) of structurally similar molecules. Synergistic interactions with FLZ labeled in green and non-interacting molecules labeled in grey. * represents FICI for 50% inhibition of *C. neoformans* all others listed are FICI for 90% inhibition.

**Figure S4.**
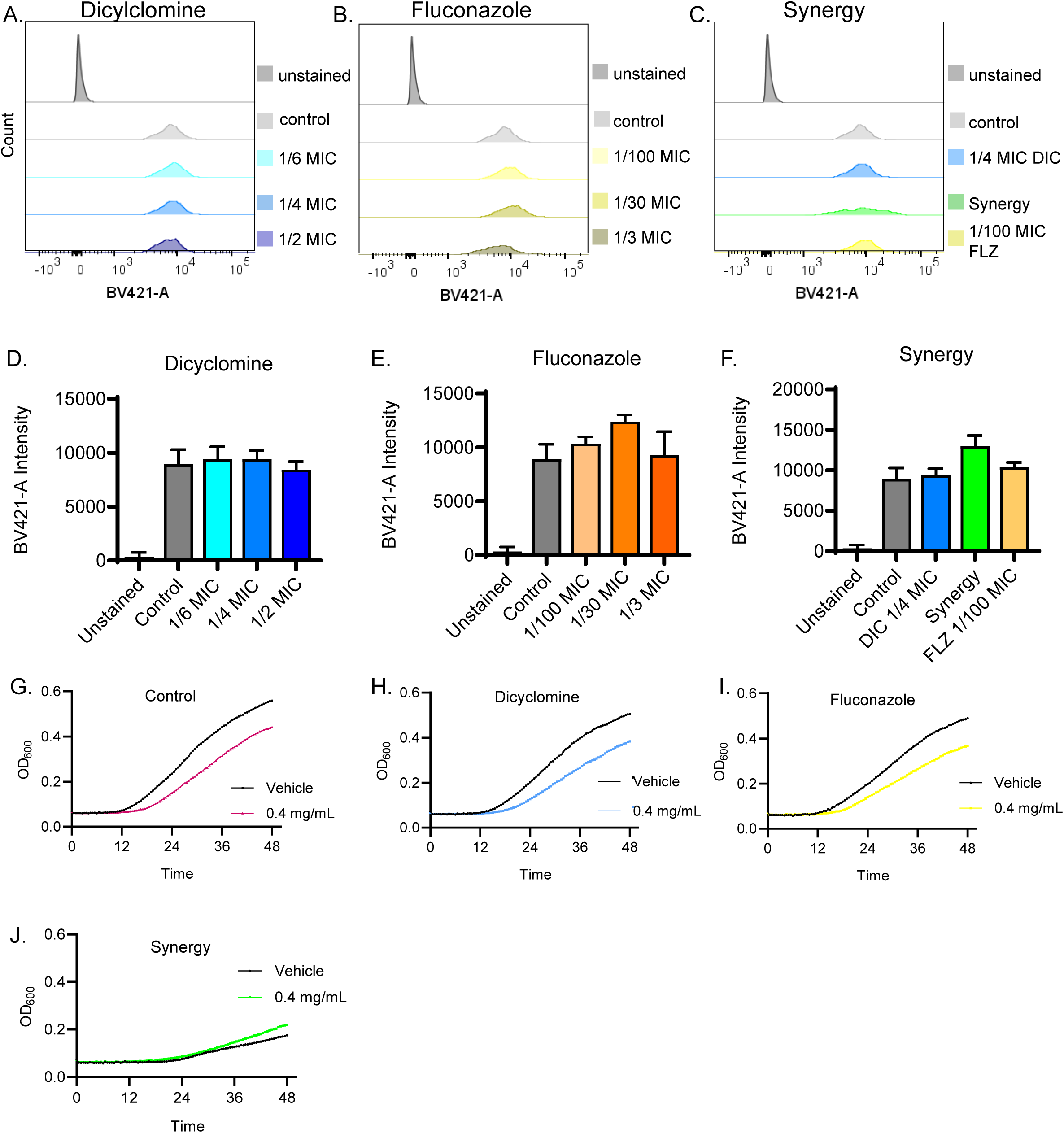
Dicyclomine additional effects on fungal chitin staining and nutrient intake. **A-C**) Representative flow cytometry of Calcofluor white staining with various concentrations of Dicyclomine (DIC) (blue), FLZ (yellow), or synergy (green). **D-F**) Quantification of average fluorescence intensity of calcofluor white staining flow cytometry. **G-J**) Growth curves of *C. neoformans* with or without 0.4 mg/mL of 5-MT after treatment with control (**G**), dicyclomine (**H**), FLZ (**I**), or Synergy (SYN) (**J**).

**Figure S5.**
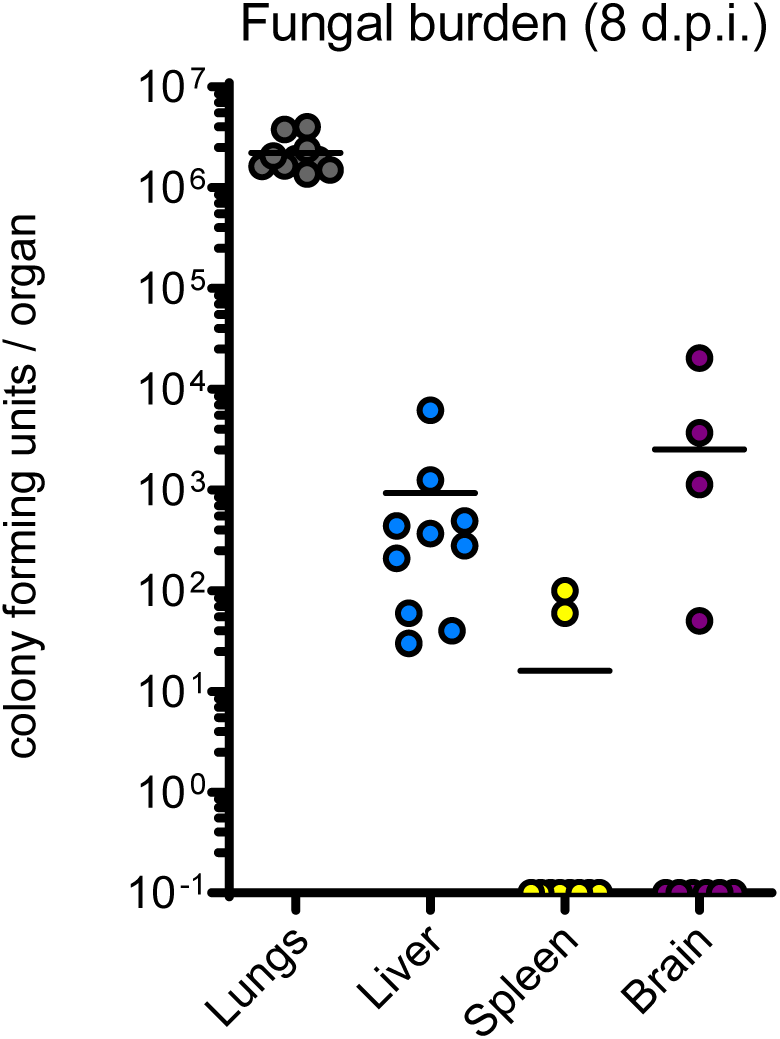
*C. neoformans* disseminates by 8 days in CD-1 outbred mice. Fungal burden of *C. neoformans* in CD-1 outbred mice at 8 days post infection.

**Table S1.** Small molecules predicted to synergize with fluconazole by O2M. Small molecules predicted to interact with FLZ. Bioactivity and Status determined by Microsource Spectrum Library. INN, International Nonproprietary Names; USAN, United States Accepted Name; BAN, British Approved Names; JAN, Japanese Adopted Name; USP, United States Pharmacopeia; NF, National Formulary.

| Small Molecule | cas # | Bioactivity | Status | Assay Used | Manufacturer |
| --- | --- | --- | --- | --- | --- |
| 1,3-Dideacetyldeoxykhivoin | N/A | N/A | Undetermined activity | not tested | N/A |
| 1,3-Dideacetylkhivoin | N/A | N/A | Undetermined activity | not tested | N/A |
| 12-Hydroxy-4,4-Bisnor-4,8,11,13-Podocarpatrien-3-One | N/A | N/A | Undetermined activity | not tested | N/A |
| 12-Methoxy-4,4-Bisnor-5alpha-8,11,13-Podocarpatrien-3-ol | N/A | N/A | Undetermined activity | not tested | N/A |
| 14-Methoxy-4,4-Bisnor-8,11,13-Podocarpatrien-3-One | N/A | N/A | Undetermined activity | not tested | N/A |
| 18alpha-Glycyrrhetic Acid | N/A | antiinflammatory | Experimental | not tested | N/A |
| 1-Hydroxy-3,6,7-trimethoxy-2,8-diprenylxanthone | 15404-76-9 | N/A | Undetermined activity | not tested | N/A |
| 1-Methylxanthine | 6136-37-4 | diuretic, adenosine antagonist | Experimental | bliss independence | VWR |
| 1-Monopalmitin | 542-44-9 | N/A | Undetermined activity | checkerboard | Sigma Chemical |
| 1-Phenylbiguanide Hydrochloride | 55-57-2 | 5HT3 receptor agonist | Experimental | bliss independence | Sigma Chemical |
| 2',2'-Bisepigallocatechin Digallate | N/A | N/A | Undetermined activity | not tested | N/A |
| 2,3,4'-Trihydroxy-4-Methoxybenzophenone | N/A | N/A | Undetermined activity | not tested | N/A |
| 2',3-Dihydroxy-4,4',6'-Trimethoxychalcone | 38186-71-9 | N/A | Undetermined activity | not tested | N/A |
| 2',4'-Dihydroxychalcone 4'-Glucoside | N/A | anthelmintic & antiulcerogenic | Experimental | not tested | N/A |
| 21-Acetoxypregnenolone | 566-78-9 | precursor in corticoid biosynthesis, derivative | Undetermined activity | bliss independence | Santa Cruz |
| 2-Hydroxy-3,4-Dimethoxybenzoic Acid | N/A | prostaglandin synthetase inhibitor | Experimental | checkerboard | Sigma Chemical |
| 2-Methoxyresorcinol | 29267-67-2 | N/A | Undetermined activity | checkerboard | Santa Cruz |
| 3,4'-Dihydroxyflavone | 14919-49-4 | N/A | Undetermined activity | checkerboard | Fisher Scientific |
| 3,4'-Dihydroxyflavone | 4143-64-0 | N/A | Undetermined activity | checkerboard | Fisher Scientific |
| 3,4-Dimethoxycinnamic acid | 2316-26-9 | N/A | Undetermined activity | checkerboard | Fisher Scientific |
| 3,4-Dimethoxydabergione | 41043-20-3 | induces dermatitis | Undetermined activity | not tested | N/A |
| 3,7-Epoxyaryophyllan-6-One | N/A | N/A | Undetermined activity | not tested | N/A |
| 3-Acetoxypregn-16-En-12,20-Dione | N/A | N/A | Undetermined activity | not tested | N/A |
| 3alpha-Acetoxydihydrodeoxygedunin | N/A | N/A | Undetermined activity | not tested | N/A |
| 3alpha-Hydroxy-4,4-Bisnor-8,11,13-Podocarpatriene | N/A | N/A | Undetermined activity | not tested | N/A |
| 3-alpha-Hydroxydeoxygedinin | N/A | N/A | Undetermined activity | not tested | N/A |
| 3-Amino-1,2,4-Triazole | 61-82-5 | catalase inhibitor | Experimental | checkerboard | Fisher Scientific |
| 3-Amino-beta-Pinene | 18172-67-3 | N/A | Undetermined activity | checkerboard | Sigma Chemical |
| 3beta-Chloroandrostanone | N/A | N/A | Undetermined activity | not tested | N/A |
| 3-Bromo-7-Nitroindazole | 74209-34-0 | NO synthetase inhibitor | Experimental | checkerboard | Fisher Scientific |
| 3-Deacetylkhivoin | N/A | N/A | Undetermined activity | not tested | N/A |
| 3-Deoxo-3beta-Acetoxydeoxydihydrogedunin | N/A | N/A | Undetermined activity | not tested | N/A |
| 3-Deoxo-3beta-Hydroxymexicanolide 16-Enol Ether | N/A | N/A | Undetermined activity | not tested | N/A |
| 3-Hydroxytyramine | 62-31-7 | dopaminergic | Experimental | not tested | N/A |
| 3-Methyl-1-Phenyl-2-Pyrazolin-5-One (MCI-186) | N/A | antioxidant, lipoxygenase inhibitor | Experimental | checkerboard | Santa Cruz |
| 4'-Methoxychalcone | 22966-19-4 | N/A | Undetermined activity | checkerboard | VWR |
| 5,7-Dichlorokynurenic Acid | 190908-40-8 | NMDA receptor antagonist (gly) | Experimental | bliss independence | Fisher Scientific |
| 5beta-12-Methoxy-4,4-Bisnor-8,11,13-Podocarpatrien-3-One | N/A | N/A | Undetermined activity | not tested | N/A |
| 5-Fluorindole-2-Carboxylic Acid | 399-76-8 | NMDA receptor antagonist (gly) | Experimental | Not tested | N/A |
| 5-Hydroxyiminoisocaryophyllene | N/A | N/A | Undetermined activity | not tested | N/A |
| 6alpha-Methylprednisolone Acetate | N/A | glucocorticoid | Experimental | not tested | N/A |
| 7,8-Dihydroxyflavone | 38183-03-8 | vascular protectant, antithrombotic | Experimental | bliss independence | Santa Cruz |
| 7-Deacetoxy-7-Oxokhivoin | 15004-51-0 | N/A | Undetermined activity | not tested | N/A |
| 7-Desacetoxy-7-Dehydrogedunin | 26927-01-5 | N/A | Undetermined activity | checkerboard | Sigma Chemical |
| 8-Cyclopentyltheophylline | 35873-49-5 | A1 adenosine agonist | Experimental | bliss independence | Fisher Scientific |
| 8-Hydroxy-15,16-Bisnor-11-labden-13-One | N/A | N/A | Undetermined activity | not tested | N/A |
| 8-Hydroxycarapinic Acid | N/A | N/A | Undetermined activity | not tested | N/A |
| 8-Iodocatechin Tetramethyl Ether | N/A | N/A | Undetermined activity | not tested | N/A |
| Acadesine | 2627-69-2 | glucose uptake stimulant; AMPK activator | Experimental | bliss independence | Cayman Chemicals |
| Acamprosate Calcium | 77337-73-6 | alcohol antagonist | USAN, INN, BAN | checkerboard | Fisher Scientific |
| Acetyltryptophan | 1218-34-4 | antidepressant | Experimental | checkerboard | Fisher Scientific |
| Acetic Acid | 585-84-2 | N/A | Undetermined activity | checkerboard | Fisher Scientific |
| Adrenaline Bitartrate | 51-42-3 | adrenergic agonist, bronchodilator, antiglaucoma agent | USP, JAN | checkerboard | Fisher Scientific |
| Aliskiren Hemifumarate | 173334-57-1 | renin inhibitor | USAN, INN | bliss independence | VWR |
| Allopurinol | 315-30-0 | antihypericemia, antigout, antilithic | USP, INN, BAN, JAN | bliss independence | Fisher Scientific |
| Alonin | 5133-19-7 | cathartic, laxative | BAN | checkerboard | Cayman Chemicals |
| Aminohydroxybutyric Acid | 924-49-2 | antiepileptic | JAN | bliss independence | Santa Cruz |
| Amiodarone Hydrochloride | 1951-25-3 | adrenergic agonist, coronary vasodilator, Ca channel blocker | USAN, INN, BAN, JAN | checkerboard | Fisher Scientific |
| Amprolium | 121-25-5 | coccidiostat | USP, INN, BAN | bliss independence | VWR |
| Anabasine Hydrochloride | 13078-04-1 | insecticide | agricultural use | not tested | N/A |
| Ancitabine Hydrochloride | 10212-25-6 | antineoplastic | INN, JAN | checkerboard | Santa Cruz |
| Andrographolide | 5508-58-7 | N/A | Undetermined activity | bliss independence | Cayman Chemicals |
| Androsta-1,4-Dien-3,17-Dione | 897-06-3 | N/A | Undetermined activity | not tested | N/A |
| Angolensic Acid, Methyl Ester | 2629-14-3 | N/A | Undetermined activity | not tested | N/A |
| Anhydrobraziliic Acid | N/A | N/A | Undetermined activity | not tested | N/A |
| Anthothecol | 10410-83-0 | N/A | Undetermined activity | not tested | N/A |
| Anthracene-9-Carboxylic Acid | 723-62-6 | Cl transport inhibitor | Experimental | checkerboard | Fisher Scientific |
| Aphyllic Acid | 642-67-1 | N/A | Undetermined activity | not tested | N/A |
| Apiin | 26544-34-3 | N/A | Undetermined activity | bliss independence | Santa Cruz |
| Arteminol | 81496-81-3 | antimalarial, antiinflammatory | INN | checkerboard | Sigma Chemical |
| Arthronioic Acid | 25556-24-5 | N/A | Undetermined activity | not tested | N/A |
| Asiatic acid | 464-92-6 | wound healing, Experimental carcinogen | Experimental | checkerboard | Sigma Chemical |
| Astaxanthin | 71772-51-5 | N/A | Undetermined activity | checkerboard | Cayman Chemicals |
| Atractyloside Potassium | 17754-44-8 | oxidative phosphorylation inhibitor | Experimental | not tested | N/A |
| Avocadyne | 34524-38-4 | antibacterial, antifungal | Experimental | not tested | N/A |
| Bambuterol Hydrochloride | 81732-46-9 | beta adrenergic agonist, bronchodilator, cholinesterase inhibitor | INN, BAN | bliss independence | Cayman Chemicals |
| Batyl Alcohol | 544-62-7 | N/A | Undetermined activity | not tested | N/A |
| Benzalkonium Chloride | 8001-54-5 | antifective (topical) | USAN, NF, INN, BAN, JAN | checkerboard | VWR |
| Berbamine Hydrochloride | 114784-59-7 | antihypertensive, skeletal muscle relaxant | Experimental | checkerboard | Cayman Chemicals |
| Berberine Chloride | 633-65-8 | antiarrhythmic, alpha2 agonist, cholinesterase, anticonvulsant, antiinflammatory, antibacterial, antifungal, antitypanosomal, antineoplastic, immunostimulant | JAN | checkerboard | Cayman Chemicals |
| Bergapten | 484-20-8 | antipsoriatic, antiinflammatory | INN | checkerboard | Santa Cruz |
| beta-Dihydrogedunol | N/A | N/A | Undetermined activity | not tested | N/A |
| beta-Escin | 6805-41-0 | membrane permeabilizer | INN | checkerboard | Fisher Scientific |
| beta-Peltatin | 518-29-6 | antineoplastic, cytotoxic | Experimental | not tested | N/A |
| Beta-Propiolactone | 57-57-8 | antiinfective | USAN, INN | not tested | N/A |
| Betonidine | 515-25-3 | N/A | Undetermined activity | checkerboard | Santa Cruz |
| Betulin | 473-98-3 | N/A | Undetermined activity | bliss independence | Cayman Chemicals |
| Bicuculline (+) | 485-49-4 | GABAa antagonist | Experimental | checkerboard | Cayman Chemicals |
| Biochanin A | 491-80-5 | phytoestrogen | Experimental | checkerboard | Cayman Chemicals |
| Biochanin A Diacetate | N/A | N/A | Undetermined activity | not tested | N/A |
| Bisabolol Acetate | N/A | N/A | Undetermined activity | not tested | N/A |
| Bismuth Subsalicylate | 14882-18-9 | antidiarrheal, antacid, antilcer | USP, JAN | checkerboard | Santa Cruz |
| Blasticidin S | 2079-00-7 | antibiotic, antifungal; LD50 (rat po) 16mg/kg | agricultural use | checkerboard | ThermoFisher |
| Bleomycin | 9041-93-4 | antineoplastic | USP, INN, BAN, JAN | checkerboard | VWR |
| Bromhexine Hydrochloride | 611-75-6 | expectorant | USAN, INN, BAN, JAN | checkerboard | VWR |
| Bromo-3-Hydroxy-4-(Succin-2-YL)-Caryolane gamma-Lactone | N/A | N/A | Undetermined activity | not tested | N/A |
| Buccladisine | 362-74-3 | vasodilator | INN, JAN | bliss independence | Cayman Chemicals |
| Bussein | 41060-14-4 | N/A | Undetermined activity | not tested | N/A |
| Cadaverine Tartrate | 462-94-2 | N/A | Undetermined activity | checkerboard | Santa Cruz |
| Calcein | 1461-15-0 | chelating agent (Ca, Mg) | Experimental | checkerboard | Cayman Chemicals |
| Caperatic Acid | 29227-64-3 | antibacterial (tuberculostatic) | Undetermined activity | not tested | N/A |
| Capobenic Acid | 21434-91-3 | antiarrhythmic | USAN, INN | not tested | N/A |
| Carbimazole | 22232-54-8 | antithyroid | INN, BAN | checkerboard | Santa Cruz |
| Caryophyllene [(+)] | 87-44-5 | N/A | Undetermined activity | checkerboard | VWR |
| Cearoin | 52811-37-7 | N/A | Undetermined activity | not tested | N/A |
| Celastrol | 34157-83-0 | antineoplastic, antiinflammatory, NO synthesis inhibitor, chaperone stimulant | Experimental | checkerboard | Cayman Chemicals |
| Cevadine | 62-59-9 | antihypertensive | Experimental | not tested | Fisher Scientific |
| Chaulmoogric Acid, Ethyl ester | 623-32-5 | antileptetic | Experimental | not tested | N/A |
| Chloroquinide Hydrochloride | 637-32-1 | antimalarial | USP-XIV, INN, BAN | checkerboard | Sigma Chemical |
| Chlorotrianisene | 569-57-3 | estrogen | USP-XIII, INN, BAN | not tested | N/A |
| Chloroxine | 773-76-2 | chelating agent | USAN | checkerboard | Fisher Scientific |
| Chlorzoxazone | 95-25-0 | muscle relaxant (skeletal) | USP, INN, BAN, JAN | checkerboard | VWR |
| Cholic Acid | 81-25-4 | N/A | Undetermined activity | checkerboard | no interaction |
| Cimicifugoside H1 | N/A | N/A | Undetermined activity | not tested | N/A |
| Cimicifugoside H2 | 161097-77-4 | N/A | Undetermined activity | not tested | N/A |
| Citrulline | 627-77-0 | N/A | Undetermined activity | bliss independence | Fisher Scientific |
| Clomipramine Hydrochloride | 17321-77-6 | antidepressant | USP, INN, BAN, JAN | checkerboard | Fisher Scientific |
| Clovanediol Diacetate | N/A | N/A | Undetermined activity | not tested | N/A |
| Colistin Sulfate | 1264-72-8 | antibacterial | USP, INN, BAN, JAN | checkerboard | Cayman Chemicals |
| Convallatoxin | 508-75-8 | cardiotonic | Experimental | checkerboard | Sigma Chemical |
| Cortisone | 53-06-5 | antiinflammatory, glucocorticoid | INN, BAN | checkerboard | Sigma Chemical |
| Cotinine | 486-56-6 | antidepressant | Experimental | checkerboard | Fisher Scientific |
| Cresol | 1319-77-3 | antiinfectant | USAN, NF, JAN | checkerboard | VWR |
| Crinamine | 639-41-8 | antihypertensive, respiratory depressant, antineoplastic | Experimental | not tested | N/A |
| Cuneatin Methyl Ether | 4253-00-3 | N/A | Undetermined activity | not tested | N/A |
| Cycloveratylene | N/A | N/A | Undetermined activity | not tested | N/A |
| Cytisine | 485-35-8 | antiinflammatory, respiratory stimulant | INN | bliss independence | Fisher Scientific |
| Dapsone | 80-08-0 | antibacterial, leprostatic, dermatitis herpetiformis suppressant | USP, INN, BAN | checkerboard | Sigma Chemical |
| Daunorubicin | 20830-81-3 | antineoplastic | USAN, INN, BAN, JAN | checkerboard | Santa Cruz |
| Decoquinat | 18507-89-6 | coccidiostat | USP, INN, BAN | bliss independence | VWR |
| Dehydro(11,12)Ursolic Acid Lactone | N/A | N/A | Undetermined activity | not tested | N/A |
| Dehydroepiandrosterone | 53-43-0 | N/A | Undetermined activity | checkerboard | Cayman Chemicals |
| Demethylnobiletin | 2174-59-6 | N/A | Undetermined activity | not tested | N/A |
| Derrusin | 14736-62-0 | N/A | Undetermined activity | not tested | N/A |
| Desipramine Hydrochloride | 58-28-6 | antidepressant | USP, INN, BAN, JAN | checkerboard | Sigma Chemical |
| Dexpanthenol | 81-13-0 | cholinergic | USP, INN, BAN | checkerboard | VWR |
| Diallyl Sulfide | 592-88-1 | antibacterial, antifungal, antineoplastic, antihypercholesterolaemic, hepatoprotectant | Experimental | checkerboard | Fisher Scientific |
| Dictamine | 484-29-7 | N/A | Undetermined activity | not tested | N/A |
| Dicyclomine Hydrochloride | 67-92-5 | anticholinergic | USP, INN, BAN, JAN | checkerboard | Sigma Chemical |
| Difucol Hexamethyl Ether | 14262-07-8 | N/A | Undetermined activity | not tested | N/A |
| Digitonin | 11024-24-1 | N/A | Undetermined activity | checkerboard | Cayman Chemicals |
| Digitoxin | 71-63-6 | inotropic, cardiotonic | USP, INN, BAN, JAN | bliss independence | Santa Cruz |
| Dihydroergotamine Mesylate | 6190-39-2 | vasoconstrictor, antimigraine | USP, INN, BAN, JAN | checkerboard | VWR |
| Dihydrofissinolate | N/A | N/A | Undetermined activity | not tested | N/A |
| Dihydrogamabogic Acid | N/A | N/A | Undetermined activity | not tested | N/A |
| Dihydrogedunin | N/A | N/A | Undetermined activity | not tested | N/A |
| Dihydrojasmonic Acid | 98674-52-3 | plant growth regulator | Experimental | checkerboard | Sigma Chemical |
| Dihydromyristicin | 607-91-0 | GSH transferase inducer | Experimental | checkerboard | Cayman Chemicals |
| Dihydrorotenone | N/A | N/A | Undetermined activity | not tested | N/A |
| Dimethylcaffeic Acid | 14737-89-4 | N/A | Undetermined activity | not tested | N/A |
| Dimethylsulfone | 67-71-0 | antiinflammatory, antiproliferative, antiparasitic | agricultural use | bliss independence | Cayman Chemicals |
| Diphenhydramine Hydrochloride | 147-24-0 | antihistaminic | USP, INN, BAN, JAN | checkerboard | Fisher Scientific |
| Disulfiram | 97-77-8 | alcohol antagonist | USP, INN, BAN, JAN | checkerboard | VWR |
| Dopamine Hydrochloride | 62-31-7 | cardiotonic, antihypotensive | USP, INN, BAN, JAN | bliss independence | VWR |
| Ebselen | 60940-34-3 | antioxidant, lipoxygenase inhibitor, inhibits oxidation of LDL | INN | checkerboard | Cayman Chemicals |
| Embelin | N/A | anthelmintic, oral contraceptive | Experimental | bliss independence | Cayman Chemicals |
| Emodic Acid | 578-45-5 | cathartic, purgative | Experimental | not tested | N/A |
| Epiandrosterone | N/A | N/A | Undetermined activity | checkerboard | Fisher Scientific |
| Epicatechin Pentaacetate | N/A | N/A | Undetermined activity | not tested | N/A |
| Epitheafavin Monogallate | N/A | N/A | Undetermined activity | not tested | N/A |
| Epoxygedunin | N/A | N/A | Undetermined activity | not tested | N/A |
| Ergosterol | 57-87-4 | N/A | Undetermined activity | bliss independence | Cayman Chemicals |
| Estradiol Benzoate | 50-50-0 | estrogen | USP-XX, INN, BAN, JAN |  | Santa Cruz |
| Estriol | 50-27-1 | estrogen | USP, INN, BAN, JAN | checkerboard | Fisher Scientific |
| Estrone | 53-16-7 | estrogen | USP, INN, BAN | checkerboard | Fisher Scientific |
| Etanidazole | 22668-01-6 | antineoplastic, hypoxic cell radiosensitizer | USAN, INN | not tested | N/A |
| Ethacrynic Acid | 58-54-8 | diuretic | USP, INN, BAN, JAN | checkerboard | Fisher Scientific |
| Eucatropine Hydrochloride | 536-93-6 | anticholinergic (ophthalmic) | USP, INN, BAN | not tested | N/A |
| Eugenol | 97-53-0 | analgesic (topical), antiseptic, antifungal | USP | checkerboard | Fisher Scientific |
| Euphorbiasteroid | 28649-59-4 | N/A | Undetermined activity | not tested | N/A |
| Filipin | 480-49-9 | antifungal | USAN, INN | checkerboard | Sigma Chemical |
| Fludrocortisone Acetate | 514-36-3 | mineralocorticoid | USP, INN, BAN, JAN | checkerboard | Sigma Chemical |
| Fluocinolone Acetonide | 67-73-2 | glucocorticoid, antiinflammatory | USP, INN, BAN, JAN | checkerboard | Fisher Scientific |
| Flurothyl | 333-36-8 | central stimulant, convulsant | USP-XXI, INN, BAN |  | VWR |
| Fumarprotocetraric Acid | 489-50-9 | N/A | Undetermined activity | not tested | N/A |
| Gallamine Triethiodide | 65-29-2 | muscle relaxant (skeletal) | USP, INN | checkerboard | Santa Cruz |
| Gemfibrozil | 25812-30-0 | antihyperlipoproteinemic | USP, INN, BAN | checkerboard | VWR |
| Gentisic Acid | 490-79-9 | analgesic, antiinflammatory | Experimental | checkerboard | Fisher Scientific |
| Geraldol | 21511-25-1 | N/A | Undetermined activity | not tested | N/A |
| Glucitol-4-Gucopyanoside | N/A | N/A | Undetermined activity | not tested | N/A |
| Glucosaminic Acid | N/A | N/A | Undetermined activity | bliss independence | Fisher Scientific |
| Glyburide | 10238-21-8 | antihyperglycemic | USP, INN, BAN, JAN | checkerboard | VWR |
| Gossypetin | 489-35-0 | N/A | Undetermined activity | checkerboard | Fisher Scientific |
| Guaiiazulene | 49-84-9 | antioxidant, inhibits lipid peroxidation inhibitor, antiinflammatory, hepatoprotectant; LD50(rat) 1550 mg/kg po | Experimental | checkerboard | VWR |
| Guaiol(-) | 489-86-1 | N/A | Undetermined activity | bliss independence | Sigma Chemical |
| Haematommic Acid | 479-25-4 | N/A | Undetermined activity | not tested | N/A |
| Harmalol Hydrochloride | 6028-07-5 | anthelmintic, narcotic agent | Experimental | bliss independence | VWR |
| Harmene | 486-84-0 | intercalating agent, sedative | Experimental | checkerboard | Santa Cruz |
| Hastatoside | 50816-24-5 | N/A | Undetermined activity | not tested | N/A |
| Hecogenin | 467-55-0 | antiinflammatory | Experimental | bliss independence | VWR |
| Hematein | 475-25-2 | N/A | Undetermined activity | not tested | N/A |
| Heptaminol Hydrochloride | 543-15-7 | vasodilator | INN, BAN | checkerboard | Sigma Chemical |
| Hetacillin Potassium | 5321-32-4 | antibacterial | USP-XIII, JAN | not tested | N/A |
| Hexetidine | 141-94-6 | antifungal | BAN | checkerboard | Santa Cruz |
| Hexylresorcinol | 136-77-6 | anthelmintic, topical antiseptic | USP, BAN | checkerboard | VWR |
| Huperzine A | 102518-79-6 | anticholinesterase, cognition enhancer | Experimental | bliss independence | Cayman Chemicals |
| Hydrocortisone Hemisuccinate | 125-04-2 | glucocorticoid | USP, JAN | bliss independence | Santa Cruz |
| Hydrocortisone Phosphate Triethylamine | 3863-59-0 | glucocorticoid | USP, INN, BAN, JAN | bliss independence | Fisher Scientific |
| Hypoxanthine | 68-94-0 | N/A | Undetermined activity | checkerboard | VWR |
| Inosine | 58-63-9 | cell function activator, cardiotonic | INN, JAN | checkerboard | VWR |
| Inositol | 87-89-8 | growth factor | NF-XII | checkerboard | Fisher Scientific |
| Iretol | N/A | N/A | Undetermined activity | not tested | N/A |
| Isopeanol | 493-33-4 | N/A | Undetermined activity | not tested | N/A |
| Isotectorigenin, 7-Methyl Ether | N/A | N/A | Undetermined activity | not tested | N/A |
| Ivermectin | 70288-86-7 | antiparasitic | USP, INN, BAN | checkerboard | VWR |
| Ketoconazole | 65277-42-1 | antifungal | USP, INN, BAN, JAN | checkerboard | Fisher Scientific |
| Khayanthone | 25279-68-9 | N/A | Undetermined activity | not tested | N/A |
| Khellin | 82-02-0 | vasodilator | INN | checkerboard | Fisher Scientific |
| Khivorin | 2524-38-1 | N/A | Undetermined activity | not tested | N/A |
| Koparin | 65048-75-1 | N/A | Undetermined activity | not tested | N/A |
| L(+/-)-Alliin | 556-27-4 | antibacterial, antioxidant | Experimental | checkerboard | Sigma Chemical |
| Lanosterol | 79-63-0 | N/A | Undetermined activity | checkerboard | Cayman Chemicals |
| Lapachol | 84-79-7 | antineoplastic, antifungal | Experimental | checkerboard | Santa Cruz |
| Lappaconitine | 32854-75-4 | analgesic, antiarrhythmic | Experimental | bliss independence | Santa Cruz |
| Larixinic Acid | 118-71-8 | N/A | Undetermined activity | checkerboard | Fisher Scientific |
| L-Deoxyvallin | 21593-77-1 | antineoplastic | Experimental | bliss independence | VWR |
| Leoidin Dimethyl Ether | N/A | N/A | Undetermined activity | not tested | N/A |
| Levalbuterol Hydrochloride | 50293-90-8 | bronchodilator, tocolytic | USAN | bliss independence | Santa Cruz |
| Linalool (+) | N/A | N/A | Undetermined activity | checkerboard | Sigma Chemical |
| Loganin | 18524-94-2 | N/A | Undetermined activity | checkerboard | Cayman Chemicals |
| Lupanyl Acid Hydrochloride | N/A | N/A | Undetermined activity | not tested | N/A |
| Lycopodine Perchlorate | N/A | N/A | Undetermined activity | not tested | N/A |
| Mangiferin | 4773-96-0 | MAO inhibitor, immunostimulant | Experimental | bliss independence | Fisher Scientific |
| Marmesin Acetate | N/A | N/A | Undetermined activity | not tested | N/A |
| Mebeverine Hydrochloride | 2753-45-9 | muscle relaxant (smooth) | USAN, INN, BAN | checkerboard | Santa Cruz |
| Melezitose | N/A | N/A | Undetermined activity | bliss independence | Fisher Scientific |
| Metameconine | N/A | N/A | Undetermined activity | not tested | N/A |
| Metergoline | 17692-51-2 | analgesic, antipyretic | INN, BAN | checkerboard | Santa Cruz |
| Metformin Hydrochloride | 1115-70-4 | antidiabetic | USAN, JAN | bliss independence | VWR |
| Methimazole | 60-56-0 | antihyperthyroid | USP, INN, BAN, JAN | checkerboard | Fisher Scientific |
| Methyl Robustone | N/A | N/A | Undetermined activity | not tested | N/A |
| Methylxanthoxylin | 23121-32-6 | N/A | Undetermined activity | not tested | N/A |
| Mexicanolide | 1915-67-9 | N/A | Undetermined activity | not tested | N/A |
| Morin | 480-16-0 | P450 and ATPase inhibitor | Experimental | bliss independence | Fisher Scientific |
| Mudulone | 481-94-7 | N/A | Undetermined activity | not tested | N/A |
| Nafcillin Sodium | 985-16-0 | antibacterial | USP, INN, BAN | checkerboard | VWR |
| N-Benzyltropan-4-ol | N/A | N/A | Undetermined activity | not tested | N/A |
| Nerol | 106-25-2 | weak estrogen receptor blocker | Experimental | checkerboard | Fisher Scientific |
| N-Methylantranilic Acid | 119-68-6 | N/A | Undetermined activity | checkerboard | Fisher Scientific |
| N-Methylisoleucine | 5125-98-8 | N/A | Undetermined activity | not tested | N/A |
| Nonic Acid | N/A | N/A | Undetermined activity | not tested | N/A |
| Nonoxynol-9 | 26027-38-3 | spermatoicide, contraceptive | USP, INN | checkerboard | Santa Cruz |
| Obliquin | N/A | N/A | Undetermined activity | not tested | N/A |
| Octopamine Hydrochloride | 104-14-3 | adrenergic agonist | INN | checkerboard | Cayman Chemicals |
| Oleanolic Acid Acetate | 4339-72-4 | N/A | Undetermined activity | not tested | N/A |
| Orsellinic Acid, Ethyl Ester | 2524-37-0 | N/A | Undetermined activity | checkerboard | Santa Cruz |
| Osthol | 484-12-8 | N/A | Undetermined activity | checkerboard | Cayman Chemicals |
| o-Veratraldehyde | 86-51-1 | N/A | Undetermined activity | not tested | N/A |
| Oxolinic Acid | 14698-29-4 | antibacterial | USAN, INN, BAN | bliss independence | Fisher Scientific |
| Pachyrrhizin | 10091-01-7 | insecticide | Experimental | not tested | N/A |
| Paenol | 552-41-0 | antibacterial | Experimental | checkerboard | Cayman Chemicals |
| Pamabrom | 606-04-2 | diuretic | USP | bliss independence | VWR |
| Pectolarin | 28978-02-1 | N/A | Undetermined activity | checkerboard | Sigma Chemical |
| Pelletierine Hydrochloride | N/A | N/A | Undetermined activity | not tested | N/A |
| Peonol Methyl Ether | 829-20-9 | N/A | Undetermined activity | not tested | N/A |
| Phenacetylamine Hydrochloride | 5468-37-1 | N/A | Undetermined activity | checkerboard | Fisher Scientific |
| Phenylbutyric Acid | 1821-12-1 | antiinflammatory, antihyperammonemic (Na salt) | Experimental | checkerboard | Fisher Scientific |
| Phloridzin | 60-81-1 | induces experimental glucosuria, antifeedant | Experimental | bliss independence | Cayman Chemicals |
| Picropodophyllotoxin | 477-47-4 | antineoplastic; 10% cytotoxicity of podophyllotoxin | Experimental | bliss independence | Sigma Chemical |
| Picropodophyllotoxin Acetate | N/A | N/A | Undetermined activity | not tested | N/A |
| Pinacidil | 85371-64-8 | K channel agonist, antihypertensive | USAN, INN | not tested | Santa Cruz |
| Pirolartine | 20069-09-4 | anti-asthma, antibronchitis | Experimental | checkerboard | Cayman Chemicals |
| Podotarin | N/A | N/A | Undetermined activity | not tested | N/A |
| Primaquine Diphosphate | 63-45-6 | antimalarial | USP, INN, BAN | checkerboard | Santa Cruz |
| Protopetrine B | 124-97-0 | antihypertensive, emetic; LD50(mouse) 0.21 mg/kg sc | INN | not tested | N/A |
| Pterixin | 13161-75-6 | muscle relaxant | Experimental | not tested | N/A |
| Punctapronin B | 93697-36-0 | N/A | Undetermined activity | not tested | N/A |
| Purpurgallin | 569-77-7 | xanthine oxidase inhibitor, antioxidant | Experimental | not tested | N/A |
| Putrescine Dihydrochloride | 333-93-7 | ornithine decarboxylase inhibitor, cell growth factor | Experimental | bliss independence | Santa Cruz |
| Pyridylidone | 77-04-3 | hypnotic, sedative | INN | bliss independence | Sigma Chemical |
| Quebrachitol | 642-38-6 | N/A | Undetermined activity | bliss independence | Santa Cruz |
| Rabeprazole Sodium | 117976-90-6 | gastric acid secretion inhibitor, cell growth factor | USAN, INN, BAN | checkerboard | VWR |
| Resveratrol | 501-36-0 | antifungal, antibacterial | Experimental | bliss independence | VWR |
| Rhetsinine | 526-43-2 | N/A | Undetermined activity | not tested | N/A |
| Rhodinyl Acetate | N/A | N/A | Undetermined activity | not tested | N/A |
| Rhodocladonic Acid | 26984-15-6 | N/A | Undetermined activity | not tested | N/A |
| Ribostamycin Sulfate | 25546-65-0 | antibacterial | INN, BAN, JAN | checkerboard | Santa Cruz |
| Risedronate Sodium Hydrate | 115436-72-1 | calcium regulator | USAN | bliss independence | Santa Cruz |
| Rotenone | 83-79-4 | acaricide, ectoparasiticide, antineoplastic, mitochondrial poison | agricultural use | checkerboard | Cayman Chemicals |
| Safrole | 9-59-7 | anesthetic (topical) and antiseptic, pediculicide | Experimental | checkerboard | Sigma Chemical |
| Salsolidine | 493-48-1 | antihypertensive | Experimental | bliss independence | Santa Cruz |
| Salsoline | 89-31-6 | antihypertensive, antihistamine | Experimental | not tested | N/A |
| Sappanone A Trimethyl Ether | N/A | N/A | Undetermined activity | not tested | N/A |
| Scopoletin | 92-61-5 | NO synthesis (inducible) inhibitor, anticoagulant | Experimental | checkerboard | VWR |
| Sertraline Hydrochloride | 79559-97-0 | antidepressant, 5HT uptake inhibitor | USAN, INN, BAN | checkerboard | VWR |
| Smilagenin | 126-18-1 | N/A | Undetermined activity | not tested | N/A |
| Solidagenone | 23534-56-7 | N/A | Undetermined activity | not tested | N/A |
| Spermidine Trihydrochloride | 334-50-9 | ornithine decarboxylase inhibitor | Experimental | bliss independence | Fisher Scientific |
| Succinylacetone | 51568-18-4 | inhibitor of heme biosynthesis | Experimental | checkerboard | Sigma Chemical |
| Sulfamethizole | 144-82-1 | antibacterial | USP, INN, BAN, JAN | checkerboard | Sigma Chemical |
| Tanshinone IIA Sulfonate Sodium | 568-72-9 | free radical scavenger | Experimental | checkerboard | Cayman Chemicals |
| Terazosin Hydrochloride | 70024-40-7 | antihypertensive | USP, INN, BAN, JAN | bliss independence | Fisher Scientific |
| Thermopsine perchlorate | 486-90-8 | N/A | Undetermined activity | not tested | N/A |
| Tilorone | 27591-69-1 | antiviral | USAN, INN | bliss independence | VWR |
| Tolfenamic Acid | 13710-19-5 | antiinflammatory, analgesia | INN, BAN, JAN | checkerboard | Cayman Chemicals |
| Toremiphen Citrate | 89778-27-8 | antineoplastic, anti-estrogen | USAN, INN, BAN | checkerboard | Santa Cruz |
| Totanol-19-Carboxylic Acid, Methyl Ester | N/A | N/A | Undetermined activity | not tested | N/A |
| Tubocurarine Chloride | 6989-98-6 | muscle relaxant (skeletal) | USP, INN, BAN, JAN | bliss independence | Santa Cruz |
| Tulobuterol | 41570-61-0 | bronchodilator, beta adrenergic agonist | INN, BAN, JAN | checkerboard | VWR |
| Tylosin Tartrate | 1405-54-5 | antibacterial | USP, INN, BAN | checkerboard | VWR |
| Tyramine | 51-67-2 | adrenergic agonist | Experimental | checkerboard | Cayman Chemicals |
| Uncarine E | 5171-37-9 | N/A | Undetermined activity | not tested | N/A |
| Ursinoic Acid | 30265-59-9 | N/A | Undetermined activity | not tested | N/A |
| Ursocholic Acid | 546-18-9 | N/A | Undetermined activity | checkerboard | Santa Cruz |
| Ursic Acid | 125-46-2 | antibacterial | Experimental | checkerboard | Santa Cruz |
| Utilin | 31218-22-1 | N/A | Undetermined activity | not tested | N/A |
| Valacyclovir Hydrochloride | 124832-27-5 | antiviral | USAN, INN, BAN | bliss independence | Fisher Scientific |
| Veratrine Sulfate | 62-59-9 | antihypertensive | Experimental | bliss independence | Sigma Chemical |
| Vincamine | 1617-90-9 | vasodilator | INN, BAN | checkerboard | Cayman Chemicals |
| Vindoline | 2182-14-1 | CNS stimulant/depressant; antihyperglycaemic | Experimental | checkerboard | Santa Cruz |
| Xylocarpus A | N/A | N/A | Undetermined activity | not tested | N/A |
| Xylomethazoline Hydrochloride | 1218-35-5 | adrenergic agonist, nasal<br>decongestant | USP, INN, BAN | checkerboard | VWR |
| Zolpidem | 82626-48-0 | sedative, hypnotic | USAN, INN, BAN | bliss independence | Fisher Scientific |

**Table S2.**
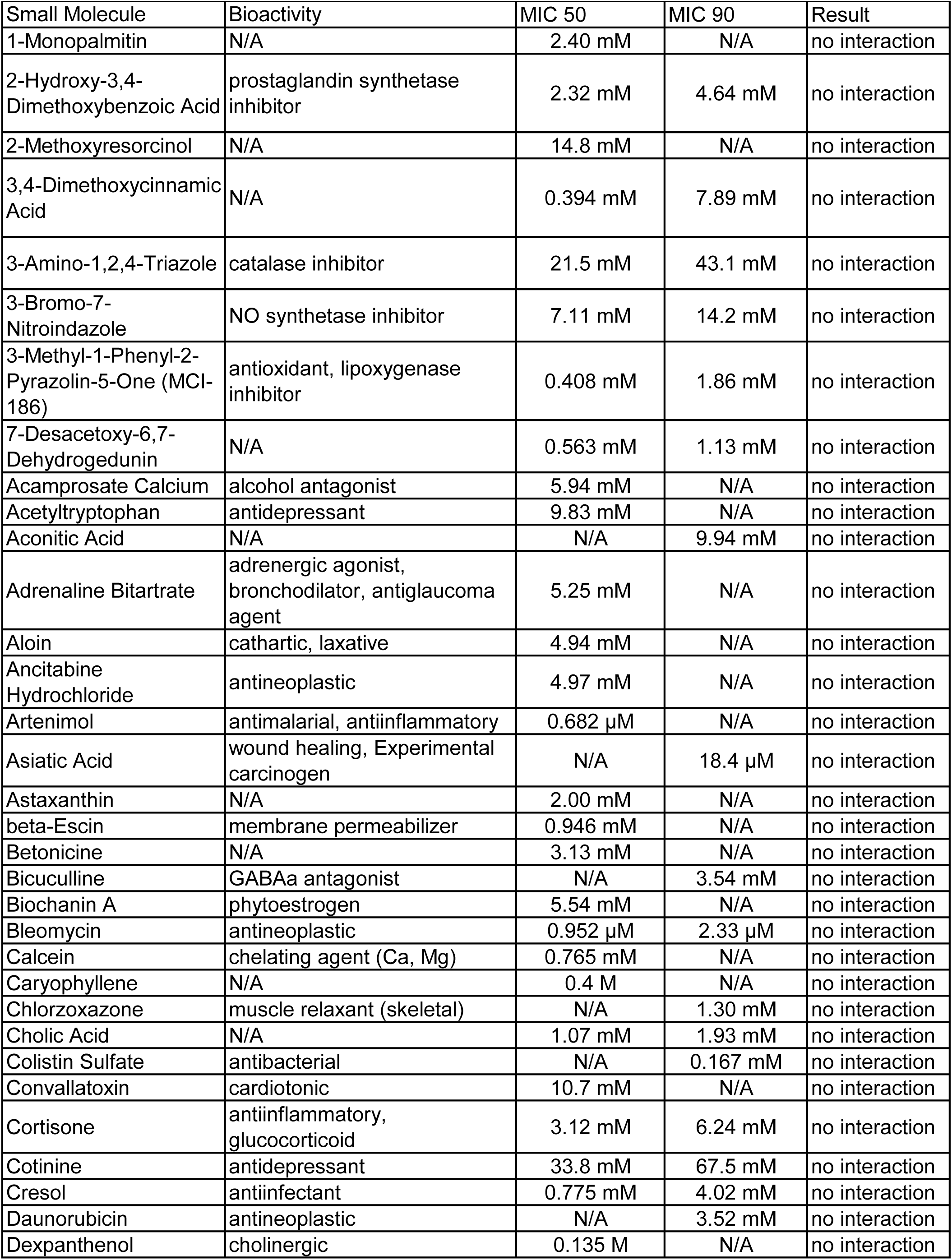
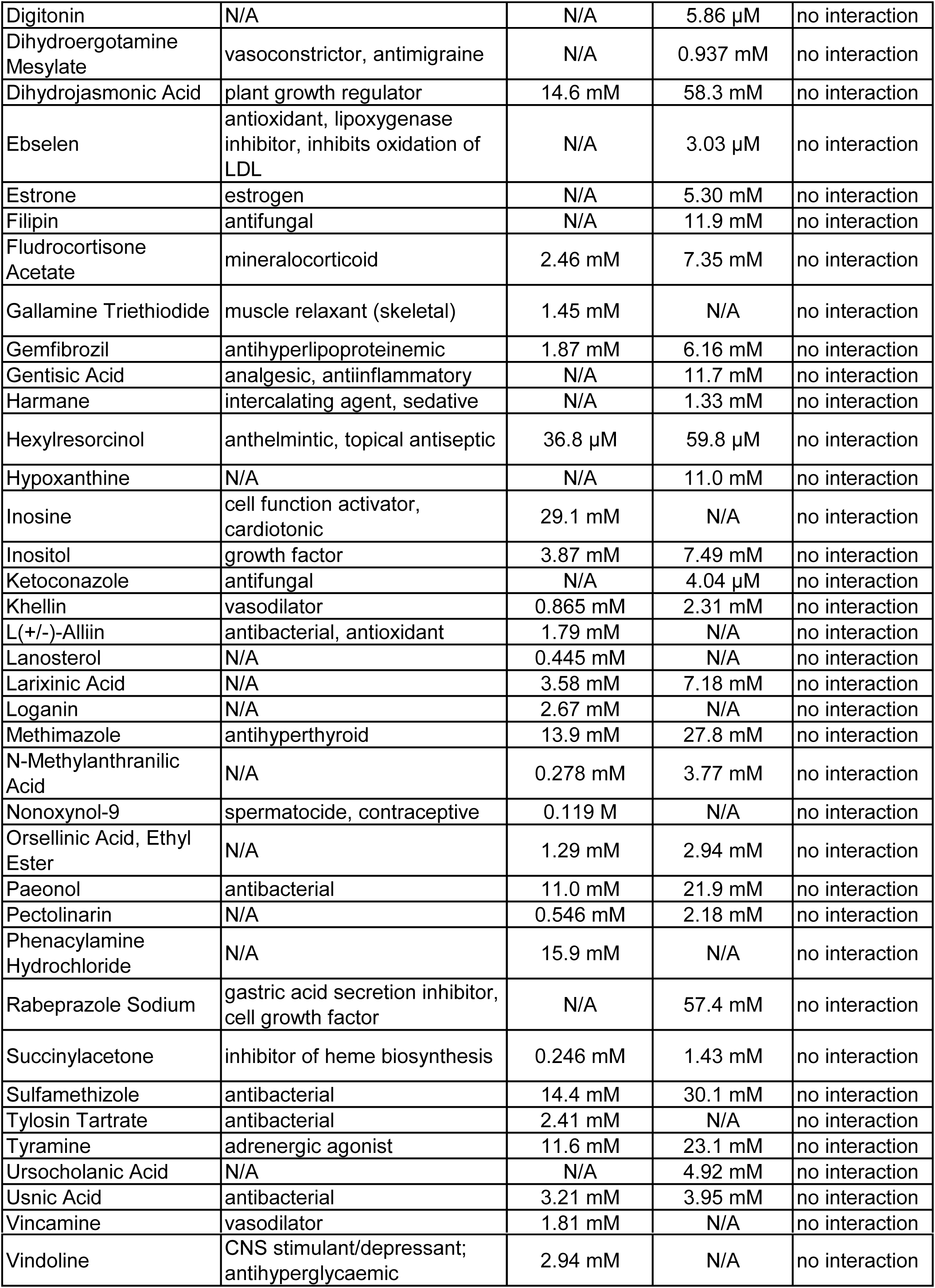
Minimum inhibitory concentrations of non-interacting molecules. Minimum inhibitory concentration for 50% inhibition (MIC 50) and 90% inhibition (MIC 90) of small molecules predicted to interact with FLZ but resulted in no interaction.

**Table S3.** Minimum inhibitory concentrations of general anti-*C. neoformans* molecules. Minimum inhibitory concentration 90% inhibition (MIC 90) for small molecules able to inhibit wild-type *C. neoformans* growth.

| Small Molecule | Bioactivity | MIC 90 |
| --- | --- | --- |
| Bithionate Sodium | anthelmintic, antiseptic | 3.70 μM |
| Cloxyquin | antibacterial, antifungal | 947 nM |
| Cycloheximide | protein synthesis inhibitor | 1.04 μM |
| Fenbendazole | anthelmintic | 37.1 nM |
| Hexachlorophene | antiinfective (topical) | 760 μM |
| Miconazole | antifungal (topical) | 507 nM |
| Oxiconazole Nitrate | antifungal | 34.9 nM |
| Phenylmercuric Acetate | antifungal | 15.9 nM |
| Pyrrithione Zinc | antibacterial, antifungal, antiseborrheic | 22.9 μM |
| Sulconazole Nitrate | antifungal | 206 nM |
| Tacrolimus | immune suppressant, antifungal | 6.44 μM |
| Thimerosal | antiinfective, preservative | 10.0 nM |
| Thiram | antifungal | 38.4 nM |

**Table S4.** Minimum inhibitory concentrations for various fungal strains/species. Minimum inhibitory concentration for 50% inhibition (MIC 50) and 90% inhibition (MIC 90) of small molecules that interacted with FLZ in various fungal species. N/A represents molecules that did not have an MIC.

|  | CM18 |  | KN99 |  | MUC 416-4 |  | MUC 402-1 |  |
| --- | --- | --- | --- | --- | --- | --- | --- | --- |
|  | MIC50 | MIC90 | MIC50 | MIC90 | MIC50 | MIC90 | MIC50 | MIC90 |
| 3-Amino-beta-Pinene | 11.7 mM | 50.1 mM | 2.43 mM | 4.51 mM | 0.293 mM | 0.879 mM | 0.535 mM | 0.967 mM |
| Benzalkonium Cl | N/A | 14.5 μM | 11.7 μM | 23.3 μM | 4.38 μM | 8.74 μM | 4.38 μM | 8.74 μM |
| Berbamine HCl | N/A | 63.5 μM | 40.2 μM | 80.5 μM | N/A | 80.5 mM | N/A | 80.5 μM |
| Bismuth Subsalicylate | 1.49 mM | N/A | 93.2 μM | N/A | 0.521 mM | 1.63 mM | 0.215 mM | 1.11 mM |
| Clomipramine HCl | 0.650 mM | 0.780 mM | 0.157 mM | 0.629 mM | N/A | 0.585 mM | N/A | 0.585 mM |
| Dehydroepiandrosterone | 1.50 mM | 12.0 mM | 1.43 mM | 11.4 mM | 0.880 mM | 8.67 mM | 3.29 mM | 8.14 mM |
| Dicyclomine HCl | N/A | 4.85 mM | 2.42 mM | 3.63 mM | N/A | 3.63 mM | N/A | 2.42 mM |
| Estriol | N/A | 24.8 mM | 12.4 mM | 18.6 mM | N/A | 17.3 mM | N/A | 19.2 mM |
| Fluocinolone Acetonide | 1.84 mM | 3.69 mM | 0.345 mM | 1.38 mM | N/A | 1.96 mM | 0.460 mM | 1.53 mM |
| Sertraline HCl | 0.209 mM | 0.313 mM | N/A | 0.214 mM | N/A | 0.209 mM | N/A | 0.209 mM |
| Xylometazoline HCl | 2.44 mM | N/A | 2.44 mM | N/A | N/A | N/A | N/A | N/A |
| Ivermectin | N/A | 1.91 mM | 0.158 mM | 1.07 mM | 0.317 mM | 0.952 mM | 1.07 mM | N/A |
| Nafcillin Sodium | 2.37 mM | N/A | 1.56 mM | N/A | 3.46 mM | N/A | 3.21 mM | N/A |
| Fluconazole | 25.0 μM | 100 μM | 24.5 μM | 98.0 μM | 18.4 μM | 49.0 μM | 12.8 μM | 0.129 mM |

| NRHc.5013.ENR |  | MUC 418-1 |  | Ftc 555-1 |  | NRHc.5027.EN.CLIN1 |  | Ftc 327-1 |  |
| --- | --- | --- | --- | --- | --- | --- | --- | --- | --- |
| MIC50 | MIC90 | MIC50 | MIC90 | MIC50 | MIC90 | MIC50 | MIC90 | MIC50 | MIC90 |
| 0.879 mM | 3.52 mM | 2.40 mM | 2.75 mM | 1.44 mM | 2.89 mM | N/A | 5.07 mM | 1.93 mM | 11.6 mM |
| N/A | 6.56 µM | N/A | 11.7 µM | 11.7 µM | 23.3 µM | 11.7 µM | 17.5 µM | 11.7 µM | 23.3 µM |
| N/A | 80.5 µM | 40.2 µM | 60.4 µM | N/A | 80.5 µM | N/A | 80.5 µM | N/A | 80.5 µM |
| 0.128 mM | 0.521 mM | 0.272 mM | 0.998 mM | 0.544 mM | 1.45 mM | 0.284 mM | 2.27 mM | 0.340 mM | 2.86 mM |
| N/A | 0.780 mM | N/A | 0.510 mM | 0.390 mM | 0.704 mM | N/A | 0.510 mM | N/A | 0.629 mM |
| 2.12 mM | 9.01 mM | 3.22 mM | 11.4 mM | 4.10 mM | 6.65 mM | 3.82 mM | 10.2 mM | 1.95 mM | 11.4 mM |
| N/A | 3.63 mM | N/A | 3.63 mM | N/A | 2.42 mM | N/A | 3.63 mM | N/A | 4.85 mM |
| 14.7 mM | 26.4 mM | N/A | 23.0 mM | N/A | 21.5 mM | 24.8 mM | N/A | N/A | N/A |
| 1.46 mM | 1.84 mM | 0.767 mM | 0.460 mM | 0.345 mM | 0.843 mM | 1.33 mM | 1.69 mM | 0.576 mM | 2.40 mM |
| N/A | 0.209 mM | N/A | 0.182 mM | N/A | 0.182 mM | N/A | 0.182 mM | 0.107 mM | 0.218 mM |
| N/A | N/A | 2.90 mM | N/A | 2.79 mM | N/A | 2.00 mM | N/A | 26.7 mM | N/A |
| 0.218 mM | N/A | 0.326 mM | 1.13 mM | 2.17 mM | N/A | 0.190 mM | 2.08 mM | 0.358 mM | 1.39 mM |
| 3.16 mM | N/A | 2.20 mM | N/A | 3.08 mM | N/A | 3.76 mM | N/A | 3.46 mM | N/A |
| 6.14 µM | 50.7 µM | 15.2 µM | 60.7 µM | 11.7 µM | 60.7 µM | 25.1 µM | 54.5 µM | 7.96 µM | 39.2 µM |

| MUC 37-1 |  | PMHC.1050.ENR.CLIN1 |  | NRHc.5011.ENR |  | *C. deuterogattii* |  | *C. auris* 1 |  |
| --- | --- | --- | --- | --- | --- | --- | --- | --- | --- |
| MIC50 | MIC90 | MIC50 | MIC90 | MIC50 | MIC90 | MIC50 | MIC90 | MIC50 | MIC90 |
| N/A | 70.8 mM | N/A | 2.90 mM | N/A | 3.65 mM | N/A | 0.967 mM | N/A | 9.69 mM |
| N/A | 5.83 µM | N/A | 11.7 µM | N/A | 23.3 µM | N/A | 5.83 µM | N/A | 46.8 µM |
| N/A | 82.5 µM | 40.2 µM | 80.5 µM | N/A | 40.2 µM | N/A | 80.5 µM | 0.161 mM | 0.322 mM |
| 0.871 mM | 1.27 mM | 1.05 mM | N/A | 0.408 mM | 0.816 mM | 0.829 mM | 2.32 mM | 0.663 mM | 2.32 mM |
| N/A | 0.532 mM | N/A | 0.629 mM | N/A | 0.629 mM | N/A | 0.629 mM | 0.629 mM | 1.26 mM |
| 3.90 mM | 10.2 mM | 3.42 mM | 9.30 mM | 8.86 mM | N/A | 0.894 mM | 1.79 mM | 4.70 mM | N/A |
| N/A | 2.42 mM | N/A | 4.85 mM | N/A | 3.63 mM | N/A | 3.63 mM | 4.85 mM | 9.66 mM |
| 24.8 mM | N/A | 24.8 mM | N/A | 6.80 mM | 26.4 mM | N/A | 24.8 mM | 15.5 mM | 18.6 mM |
| 0.734 mM | 1.38 mM | 1.46 mM | 2.08 mM | 0.468 mM | 1.15 mM | 0.576 mM | 1.38 mM | N/A | 0.756 mM |
| N/A | 0.200 mM | N/A | 0.214 mM | N/A | 0.214 mM | N/A | 0.214 mM | 0.214 mM | 0.429 mM |
| 2.67 mM | N/A | 2.73 mM | N/A | 2.67 mM | N/A | 3.77 mM | N/A | N/A | N/A |
| 0.715 mM | N/A | 0.221 mM | N/A | 0.156 mM | 0.207 mM | 0.149 mM | N/A | 0.198 mM | N/A |
| 2.87 mM | N/A | 3.60 mM | N/A | 2.88 mM | N/A | 1.45 mM | N/A | 3.46 mM | N/A |
| 52.6 µM | 0.111 mM | 24.0 µM | 90.8 µM | 21.4 µM | 85.9 µM | 40.8 µM | 81.6 µM | 1.33 mM | 4.00 mM |

| *C. auris* 2 |  | *C. albicans* |  | *C. glabrata* |  |
| --- | --- | --- | --- | --- | --- |
| MIC50 | MIC90 | MIC50 | MIC90 | MIC50 | MIC90 |
| 4.83 mM | 9.69 mM | N/A | 0.150 M | N/A | 0.150 M |
| 31.1 µM | 62.3 µM | N/A | 7.29 µM | 5.38 µM | 10.8 µM |
| N/A | 0.322 mM | N/A | 1.01 mM | 0.126 mM | 0.236 mM |
| 1.57 mM | N/A | 0.497 mM | 1.99 mM | 0.491 mM | 1.48 mM |
| N/A | 1.26 mM | N/A | 0.555 mM | 0.660 mM | 1.32 mM |
| 2.86 mM | 11.4 mM | N/A | 11.4 mM | 0.888 mM | 2.13 mM |
| 4.85 mM | 9.66 mM | N/A | 4.85 mM | 4.74 mM | 7.15 mM |
| 6.98 mM | 24.8 mM | N/A | 24.8 mM | 6.13 mM | 12.2 mM |
| N/A | 0.460 mM | 3.69 mM | N/A | N/A | 3.96 mM |
| 0.214 mM | 0.429 mM | N/A | 0.858 mM | 1.41 mM | N/A |
| N/A | N/A | 2.44 mM | N/A | 3.13 mM | N/A |
| 0.119 mM | N/A | 1.54 mM | N/A | 0.296 mM | 0.592 mM |
| 3.16 mM | N/A | 3.16 mM | N/A | 2.43 mM | N/A |
| 1.16 mM | 4.00 mM | 6.25 µM | 12.5 µM | 0.564 mM | 1.11 mM |

**Table S4.** Gene deletion mutants resistant to Dicyclomine. Gene knockouts in KN99 resistant to dicyclomine at 1.65 mg/mL.

|  |  |
| --- | --- |
| **Table S5.** |  |
| CNAG | Gene |
| CNAG_00122 | alpha-amylase |
| CNAG_00185 | hypothetical protein |
| CNAG_00186 | mitotic spindle organizing protein 1 |
| CNAG_00651 | ATP-binding cassette, subfamily D (ALD), peroxisomal long-chain fatty acid import protein |
| CNAG_00696 | alpha-mannosidase |
| CNAG_00723 | hypothetical protein |
| CNAG_00727 | mitochondrial protein with role in iron accumulation |
| CNAG_00872 | ATP-dependent DNA helicase |
| CNAG_00875 | rab family protein |
| CNAG_01002 | Hypothetical protein |
| CNAG_01192 | Hypothetical protein |
| CNAG_01211 | Hypothetical protein |
| CNAG_01212 | Hypothetical protein |
| CNAG_01247 | Hypothetical protein |
| CNAG_01248 | Vacuole morphology and inheritance protein 14 |
| CNAG_01249 | Hypothetical protein |
| CNAG_01252 | Thiosulfate/3-mercaptopyruvate sulfurtransferase |
| CNAG_01256 | Hypothetical protein |
| CNAG_01585 | Hypothetical protein |
| CNAG_01975 | Helicase |
| CNAG_02016 | hypothetical protein |
| CNAG_02057 | URE6; urease accessory protein |
| CNAG_02058 | Hypothetical protein |
| CNAG_02108 | GTPase activating protein |
| CNAG_02409 | LIV5; protein of unknown function |
| CNAG_02416 | hypothetical protein |
| CNAG_02715 | alpha-1,6-mannosyltransferase |
| CNAG_02733 | monosaccharide transporter |
| CNAG_03041 | hypothetical protein |
| CNAG_03043 | hypothetical protein, variant 1 |
| CNAG_03056 | hypothetical protein |
| CNAG_03460 | phosphoglycerate dehydrogenase |
| CNAG_03523 | solute carrier family25, member 38 |
| CNAG_03582 | RIM20; pH-response regulator protein palA/RIM20 |
| CNAG_03770 | hypothetical protein |
| CNAG_03828 | aromatic amino acid aminotransferase I |
| CNAG_03831 | hypothetical protein |
| CNAG_03871 | phosphotyrosine protein phosphatase |
| CNAG_03905 | hypothetical protein |
| CNAG_03908 | wd-repeat protein |
| CNAG_03938 | nonmating-type specific pheromone GPCR; CPR2 |
| CNAG_03945 | hypothetical protein |
| CNAG_03989 | hypothetical protein |
| CNAG_04155 | small nuclear ribonucleoprotein E |
| CNAG_04160 | hypothetical protein, variant |
| CNAG_04199 | hypothetical protein |
| CNAG_04203 | hypothetical protein |
| CNAG_04311 | charged multivesicular body protein 7 |
| CNAG_04313 | NADPH2 dehydrogenase |
| CNAG_04618 | Hypothetical protein, variant |
| CNAG_05065 | hypothetical protein |
| CNAG_05075 | solute carrier family 20 (sodium-dependent phosphate transporter) |
| CNAG_05838 | RGD1; rho GTPase activating protein homolog |
| CNAG_06225 | hypothetical protein |

